# Kinetic Lipidomics: Quantifying *in vivo* changes in lipid metabolism using metabolic labeling

**DOI:** 10.64898/2026.06.29.735310

**Authors:** Coleman O. Nielsen, Russell L. Denton, Chad R. Quilling, Kenneth L. Virgin, Kyle J. Cutler, Spencer S. Gates, Martin D Sorensen, Mitchel F. Poulson, Tayler C. Hilton, Benjamin K. Driggs, Zabdi Hernandez, Parker M. Snedaker, Bradley C. Naylor, Mark K. Transtrum, John C. Price

## Abstract

Lipid metabolism reflects the dynamic balance between metabolic turnover and concentration. Kinetic mass spectrometry (MS) enables direct quantification of molecular turnover *in vivo.* Previous work has shown that MS-based kinetic proteomics has provided powerful insights into proteome regulation. Analogous lipidome-wide kinetic measurements remain limited by challenges in defining molecule-specific labeling behavior. Here, we extend kinetic MS to untargeted lipidomics.

Isotope labeling with deuterated water (^2^H_2_O) is commonly used for monitoring turnover of palmitate and other select lipids by measuring labeling of stable C–H positions with deuterium (^2^H). Here, we extend the deuterium-incorporation model underlying these targeted lipid turnover assays to support untargeted analysis of all detectable lipids. This allows us to empirically quantify the effective fraction of endogenous synthesis (*A*_syn_) and the turnover rate (*k*) across hundreds of lipid species simultaneously.

One central barrier to lipidome-wide kinetic modeling is determining the endogenous number of deuterium-labeling sites for each molecule (*n_L_*) which is required to estimate *A_syn_* and *k* accurately. The *n_L_* value is an essential component of biological kinetic assays. In kinetic proteomics, curated amino acid *n_L_* libraries enable peptide-level modeling by summing sequence-specific labeling-site values, but comparable resources are lacking for lipids and may not generalize across metabolic states or non-mammalian systems. Yet, gaps remain for lipids and for amino acids in modified metabolic conditions or non-mammalian biologies. Here, we empirically determine lipid *n_L_* values and validate the process with peptides against an *n_L_* library.

To evaluate this strategy in a biologically relevant setting, we applied it to brain tissue from transgenic mice expressing human ApoE isoforms, where altered lipid transport and metabolism are implicated in Alzheimer’s disease risk. These data validate the method in a clinically relevant context and suggest that genotype-dependent metabolism can alter empirically determined lipid n_L_ values.

## Introduction

Lipid homeostasis in humans is regulated by balancing intestinal uptake, synthesis and degradation, transport, and excretion (Fig 1A). Loss of lipid homeostasis is implicated in various human diseases, such as cancer ^1–3^, cardiovascular disease ^4, 5^, diabetes, dysfunction-associated steatotic liver disease^6,7^, and Alzheimer’s disease ^8^. Monitoring changes in steady-state lipid abundance is not sufficient to distinguish between contributions in this dynamic system.

**Figure 1:**
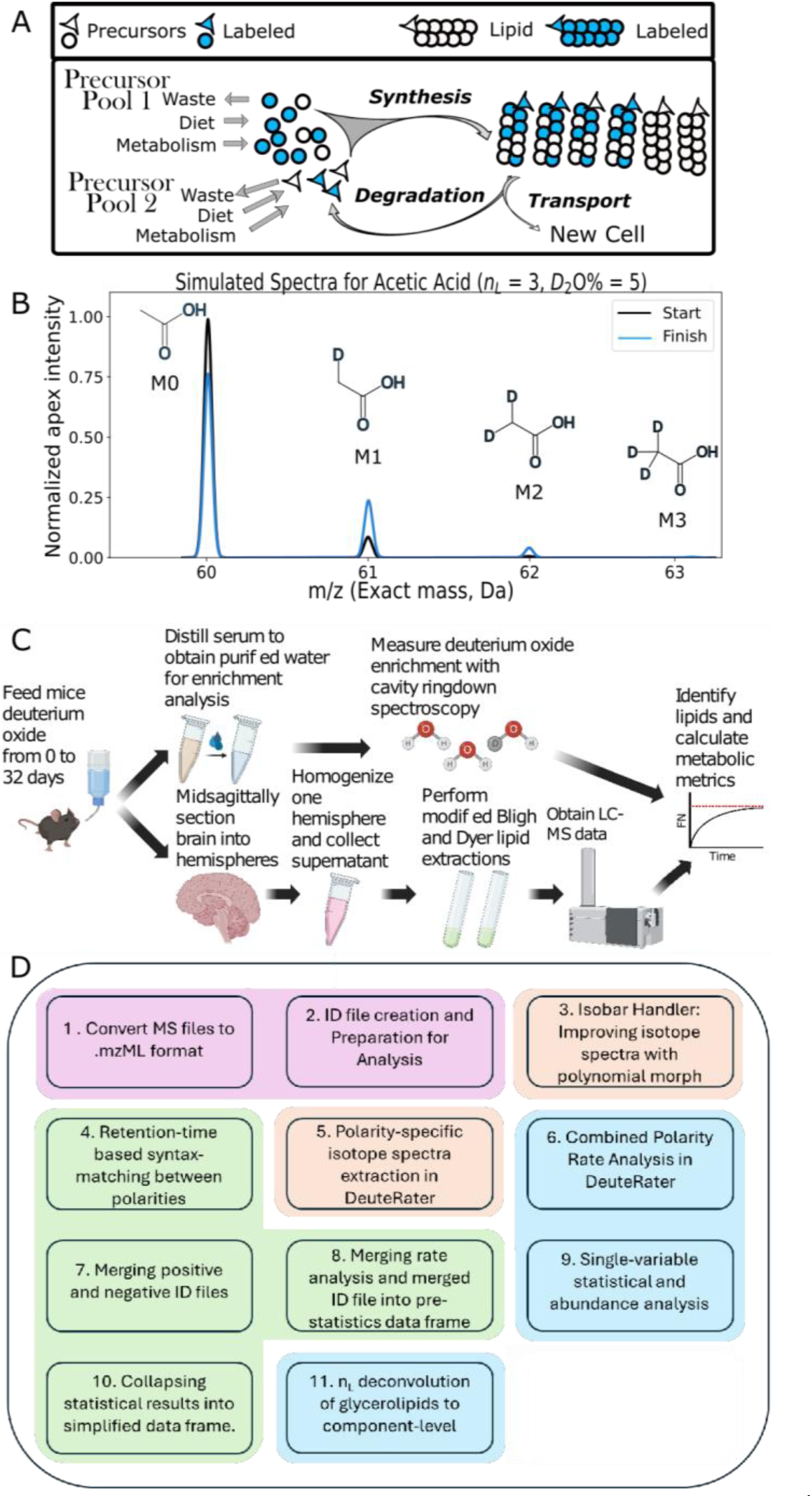
Theoretical and Experimental Summary. (A) Homeostasis model (B) Simulated isotope distributions for acetic acid (unlabeled, black and labeled, blue) (C) Experimental workflow (D) Kinetic Lipidomics Workflow using DeuteRater v7 pink= msconvert, orange=mzML handling, green=data reformatting, blue=analysis steps

Kinetic analysis of metabolic pathways directly measures the change in synthesis or degradation providing mechanistic insight ^9, 10^. Kinetic analysis can be accomplished by quantification of molecular replacement *in vivo* using the incorporation of stable isotopes into newly synthesized molecules ^11, 12^. MS-based kinetic proteomics workflows by our group ^13–15^ and others^16–18^ can measure thousands of protein turnover rates in complex mixtures. However, analogous lipidome-wide kinetic measurements remain less developed because lipid-specific labeling parameters are not generally known.

Deuterated water (²H₂O) labeling is particularly well-suited for in vivo studies in model organisms ^14, 19, 20^ and in humans ^12, 21, 22^. A key advantage of deuterium (²H) from body water is the fast distribution across the entire organism and metabolic incorporation into stable carbon–hydrogen (C–H) bonds during precursor synthesis (**Fig 1A**). The isotope labeling increases the relative abundance of heavier neutromers (chemically identical species that differ only in neutron number^23^, **Fig 1B**). While body-water enrichment (*p*) and elemental composition can be measured or inferred directly, the number of metabolically incorporated deuterium positions (*n_L_*, Fig 1B) is molecule-specific and often unknown.

Kinetic proteomics succeeds largely because amino acid *n_L_* values have been extensively characterized^14, 24, 25^ and can be summed across peptide sequences. For example, in mice amino acid *n_L_* values have been measured using tritium labeling ^24^, best-fit isotope–distribution of peptides measured at maximal labeling ^26^, and protein hydrolysis followed by gas-chromatography-mass-spectrometry^14, 27^. The tritium-derived mouse *n_L_* values agree well with the experimentally observed asymptotic spectra of human peptides^12^ indicating that human amino acid *n_L_* values are similar across mammalian systems^14, 19, 20^. Several tools for proteomics rely on this predictability such as DeuteRater ^13^, D2Ome^17^, and Riana^20^. During deuterated water labeling most biosynthesized molecules in these samples are also labeled at C-H bonds, but extending this principle to untargeted lipidomics is challenging because the *n_L_* is not known for most lipid components ^28^.

Consequently, ²H-based lipid turnover analysis has largely remained targeted, with palmitate serving as the canonical example^29^. Although in situ estimation has been explored for a limited number of lipids ^28^, lipidome-wide application requires simultaneous determination of *n_L_*, synthesis fraction, and turnover rate across many molecular species. Here we demonstrate this can be accomplished by tracking time-dependent isotope distribution changes ^10, 11^ for each molecule during continuous isotope administration.

Because many lipid species share recurring structural constituents, lipidome-wide measurements also create an opportunity to infer labeling behavior below the intact-molecule level similar to previous treatments of proteins^25^. By comparing isotope-distribution changes across structurally related lipids, n_L_ contributions from fatty acyl chains, headgroups, and glycerol backbones can be demixed and assigned component-level *n_L_* values. This enables inference of pathway-specific synthesis and dietary contributions at the constituent level, information that cannot be directly obtained from individual intact lipid species.

In addition to *n_L_*, lipid kinetic analysis requires explicit estimation of the endogenously synthesized fraction (A_syn_) because not all lipid molecules in a measured pool are necessarily replaced through endogenous synthesis during labeling. A_syn_ represents the partitioning of biologically active lipid within the total pool in vivo. Mathematically, it corresponds to the asymptotic plateau reached when the fraction newly synthesized, FN, is plotted over time. In kinetic proteomics applications, A_syn_ is often approximated as 1 because of the large number of rapidly metabolized nonessential amino acids and well mixed pools of protein. For lipids, compartmentalized metabolism can produce species-specific asymptotic plateaus below 1, due to dietary influx, or stored lipid^28^ which vary due to lifestyle and disease^8^. The kinetic lipidomics approach directly measures changes in these contributors by quantifying the A_syn_^28^.

To test whether this framework can resolve biologically meaningful differences in lipid metabolism, we applied it to brain tissue from transgenic mice expressing human ApoE isoforms (**Fig 1C**). ApoE is a lipid-transport protein found in the serum and cerebrospinal fluid (CSF). The ApoE4 is the strongest genetic risk factor for late-onset Alzheimer’s disease (AD)^30^. Because ApoE genotype alters lipid transport and is linked to AD risk, ApoE-targeted replacement mice provide a biologically relevant system for evaluating whether kinetic lipidomics can detect genotype-dependent remodeling of lipid metabolism. Together, this work establishes an untargeted ²H₂O-based kinetic lipidomics workflow, validates empirical estimation of lipid n_L_ and A_syn_, applies the method to ApoE-dependent brain lipid metabolism, and provides a graphical user interface (GUI)-based tool to support broader adoption (**Fig 1D**).

## Methods

### Mouse Models and Metabolic labeling

All experiments were performed under the approval of the Institutional Animal Care and Use Committees of Brigham Young University in conformity with the Public Health Service Policy on Humane Care and Use of Laboratory Animals. Transgenic mouse models (ApoE2: 004632 - B6.Cg-Tg(GFAP-APOE_i2)14Hol APOEtm1Unc/J, ApoE3: 004633 - B6.Cg-Tg(GFAP-APOE_i3)37Hol APOEtm1Unc/J, ApoE4: 004631 - B6.Cg-Tg(GFAP-APOE_i4)1Hol APOEtm1Unc/J, ApoE-null: B6.129P2-APOEtm1Unc/J) were purchased from Jackson Laboratories and maintained at BYU less than 8 generations from the purchased founders. Metabolic labeling was initiated by an intraperitoneal injection of D_2_O saline. Mice were then fed 8% D_2_O in their drinking water for 0, 0.25, 1, 4, 16, and 32 days at which times their tissues were collected, flash frozen on a block of dry ice and stored at -80°C until they were prepared for MS analysis. (**Fig 1**).

### Measurement of ²H enrichment

Similar to previous reports ^31^, blood isotope enrichments were measured in duplicate for all mice and averages are reported (**Table S1**). Briefly, 100 µL aliquots of each sample were placed in the caps of inverted, sealed conical screw-capped vials and distilled overnight at 80 °C. The distillate and a standard curve were individually diluted 1:300 in 18.2 Megohm water. The molar percent excess (MPE) ²H was measured on a cavity ring-down water isotope analyzer (Los Gatos Research [LGR], Los Gatos, CA, USA) similar to the published method ^32^.

### Lipid extraction and mass spectrometry

Lipids were extracted from homogenized brain tissues with a modified Bligh and Dyer technique as previously described ^2, 33^. Briefly, tissues were homogenized using a FastPrep 24 bead beater in tissue lysis buffer (0.1M Tris-HCl at pH 7.2) containing fresh Halt protease and phosphatase inhibitor cocktail (ThermoFisher). For normalization purposes an aliquot of the tissue homogenate was diluted 1/1 in 10% sodium dodecyl sulfate (SDS) for protein quantitation using bicinchoninic acid (BCA, Pierce). Under an argon atmosphere, lipids were extracted from homogenate aliquots containing 400 ug of protein under argon using 1mL of chloroform/methanol/isopropanol (3:1:1.25, v/v/v) containing internal standards (**Table S2**). The organic lipid-containing layer was then concentrated under reduced pressure at room temperature. This sample was stored under argon at - 80°C until LC-MS analysis. All samples were run in MS^1^-only mode to collect the highest quality isotope spectra. Timepoints 0, 0.4, and 1 Days of labeling were also run in data-dependent acquisition (DDA) MS^2^ mode to provide block-specific identification and MS^2^ based RT mapping against the PQC library.

### Lipid identification

Raw data files as well as details of the mass spectrometric methods and data acquisition are provided through Metabolomics Workbench (**DataTrackID7141**). Pooled quality control (PQC) samples, prepared from unlabeled lipid extracts of three mice, were separated by C18 reverse-phase high-performance liquid chromatography (HPLC) and used to collect collision-induced dissociation spectra (MS²) for molecular identification. Each genotype-specific PQC was analyzed using iterative non-targeted MS² on the Agilent 6560-A instrument, in which previously fragmented precursor ions are automatically excluded in subsequent runs to progressively expand lipid coverage. The resulting MS² spectra were converted to the open .mzML format using ProteoWizard’s MSConvert^34^ v3.0.2 and analyzed in MS-DIAL^35^ v5.5.25 to assign lipid identities, chemical formulae, and reference chromatographic elution times (retention times, RT).

### Accurate per-file LC-MS retention times via Legendre-polynomial warping

Quantifying isotope labeling over time requires accurate extraction of isotopic envelopes for each lipid across all MS¹ files. However, even modest retention-time (RT) drift (∼20 seconds in this study, dependent on closeness of isobar elution times) between runs produces inconsistent isotope envelope sampling when a single RT window per analyte is used for downstream kinetic modeling in DeuteRater^13^ (**Fig S1A**). This is because, while DeuteRater filters out instrument scans with internal envelope minima (New **Fig S2**), if two envelopes of the same mass are collected within the same extraction window, they are treated as the same molecule. This leads to mis-sampled envelopes, increased noise in turnover fits, and erroneous isotope-ratio measurements. To correct for this drift, the mass-to-charge (m/z) values taken from above-mentioned MS-DIAL lipid identity data frame were used to extract M, M+1, and M+2 extracted-ion chromatograms (EICs) for every analyte across all MS¹ runs (**Fig S1A**). Each EIC was subsequently morphed into the retention-time space of a designated reference file using a Legendre-polynomial warp. Each analyte is fit per file using a Legendre polynomial (default degree = 5), optimized using SciPy’s differential_evolution solver (v1.17) applied to smoothed, down sampled (for computational efficiency) EICs. This produces a forward warp (Pol1) that maps each file’s RT axis onto the reference frame. After applying Pol1, the morphed EICs for all files were summed and normalized to generate a stabilized reference trace (**Fig S1B**). Peaks were identified on this summed trace and retained only if they met multiple criteria: 1. be within ±30 s of the MS-DIAL-identified RT from the reference file, 2. ≥ 5% of the global maximum intensity, and 3. satisfying a signal-to-noise requirement of 1.5-times the local baseline estimate. Each accepted peak center was then back-transformed to each MS¹ file using a reverse warp (Pol2) specific to each MS1 data file. A narrow, file-specific search window was used to snap each back-transformed peak to the true local maximum of the native EIC. This produced file-specific retention times to guide the isotopic data extraction of every lipid. This RT-matching framework improved isotopic-envelope fidelity across time-course datasets and increased the number of lipids passing downstream DeuteRater quality-control filters (New **Fig S3**)^13^.

### Quantification of isotope labeling in sample-specific MS1 file

Isotopic envelopes for each lipid in the ID file were retrieved from MS¹ spectra in the corresponding .mzML files using the adduct-specific m/z and sample-specific Legendre-warped RTs. In reverse-phase chromatography, differences in unsaturation are often insufficient to produce baseline separation between closely related lipid species. Each additional double bond decreases the molecular mass by approximately 2 Da, which shifts the isotope envelope of the more unsaturated lipid into the mass range occupied by lower-order isotopes of the less unsaturated lipid. As a result, lipid species that differ by a single desaturation can produce overlapping isotope envelopes when they coelute, making reliable kinetic analysis impossible for the convoluted features. (**Fig S2A**). Here, a time-specific isotope envelope for each identified lipid was extracted from each scan of the mzML using DeuteRater as previously described ^13^. Quality scans were identified using several criteria: 1. each scan contained at least three correctly spaced isotope peaks (M₀, M₁, M₂), 2. there cannot be an internal minimum, and 3. the S/N ratio must exceed 10% of the most intense scan. An internal minimum occurs when the MS1 spectra contains overlapping spectra as illustrated in **Fig S2**. A continuous sequence of 10–60 scans were summed together for the experimental master isotopic envelope for the adduct in the specific sample (**Fig S3**).

Within DeuteRater v7, spectral quality was further ensured by enforcing RT alignment among heavy neutromers and adducts, defined here as different ion-pair forms of the same molecule with different charge carriers (e.g., H⁺ and Na⁺; **Fig. S3**), consistent with published algorithms. Neutromers of the same molecule are nearly chemically identical and therefore coelute in reverse-phase HPLC, while adducts form during electrospray ionization after chromatographic separation and therefore provide independent confirmation of the same chromatographic feature. For each molecule and its associated adducts, the intensities of three neutromers (M₀, M₁, and M₂) were compared across the retention-time window. Candidate peaks were scored using weighted quality-control metrics that prioritized coelution of neutromers and adducts, agreement with expected isotope spacing, and sufficient signal relative to local baseline noise. Molecules whose neutromer or adduct peaks failed to align in RT, exhibited internal minima, or showed low signal-to-noise were excluded from further analysis because these cases indicate unreliable identifications, insufficient signal, or spectral convolution with coeluting species. The highest-scoring candidate peak was then used to represent each molecule during extraction. Isotopic envelopes for each lipid in the ID file for each adduct-specific m/z showed higher accuracy using the sample-specific Legendre-warped RTs (**Fig S4**).

### The hydrogen isotope fitting model

The data analysis used a binomial distribution as a model of the hydrogen-only form of each molecule containing *n_L_* additional neutrons as a time-dependent mixture of two components: (i) residual preexisting molecules and (ii) newly synthesized molecules that incorporate deuterium. Hydrogen-only isotopic spectra were generated by representing the observed isotope envelope as the linear convolution of a hydrogen-only distribution with the distribution of the other elements (Carbon, Nitrogen, Oxygen, Phosphorous and Sulphur). The hydrogen-only distribution was recovered by solving a non-negative least squares (NNLS) inverse problem, a standard approach for deconvolving overlapping isotope patterns ^31, 36^. This method enforces physical nonnegativity and normalizes recovered spectra to sum to one.

NNLS optimization was implemented using bounded-variable least-squares (BVLS) routines from the SciPy scientific computing library (v1.17), applied directly to the inferred hydrogen-only distribution for each spectrum. This allows the classical mass isotopomer distribution analysis (MIDA^10^) to be applied to all timepoints of a kinetics experiment simultaneously (kinetic MIDA (kMIDA), **Equation 1**).

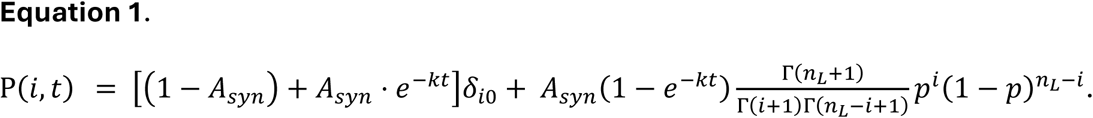

In **Equation 1**, *P(i,t)* represents the hydrogen-only distribution containing exactly *i* incorporated deuterons with a molar enrichment (*p*) at labeling time, *t*. We model *P(i,t)* as a time-dependent (*t*) mixture of a residual unlabeled (“old”) component and a labeled

(“new”) component containing a set number of metabolically incorporated labeling sites (*n_L_*). The conversion from new to old is assumed to be a first-order turnover rate (*k*) with a labeling plateau, *A_syn_*, representing the fraction of the observable pool accessible to synthetic processes over the duration of the experiment. The labeling probability *p* represents both the environmental deuterium concentration and the probability of deuterium incorporation for a stable hydrogen-incorporation site.

The Kronecker delta, δ_i0_, serves as a mathematical switch (1, when i == 0; else 0), allowing the model to also predict the unlabeled (M0) peak, which reflects changes in the pre-enrichment (“old”) molecule population. Equation 1 is expressed using the Gamma function (Γ), which generalizes factorial terms in the binomial formulation and thereby allows the effective labeling parameter *n_L_* to vary continuously. This formulation eliminates the binomial theorem’s restriction to integer *n_L_* values and accommodates effective fractional *n_L_* values arising from multiple enzymatic pathways contributing to the same molecular pool.

Empirical hydrogen-only isotope distributions were fit to Equation 1 using SciPy’s optimization routine (v1.17) with weighted least squares. Fits incorporated the first three neutromers for molecules below and four neutromers when above 1400 Da. Individual neutromer contributions were weighted by signal-to-noise so that low-confidence measurements contributed less to parameter estimation. Fraction new from same-day time points were averaged before fitting to reduce technical variability. To maintain consistency with prior versions of DeuteRater, parameter uncertainty was estimated from the covariance matrix returned by SciPy’s curve_fit function (v1.17). These covariance estimates were then used in a Wald hypothesis-testing framework to evaluate kinetic differences between genotypes. Finally, the fitted n_L_ values were used to generate theoretical isotope spectra, which provided the basis for the spectral quality filtering described below.

### Filtering on the fraction new (FN) and spectral quality for each measurement

Following annotation of each lipid ID with its *n_L_*, theoretical mass spectra were generated from the corresponding chemical formula using the eMass algorithm ^37^ as per previous versions of DeuteRater. This allows a spectral quality filter to be imposed on each lipid measurement prior to a final recalculation of kinetic constants. This is accomplished by calculating an unlabeled spectrum (**Fig 1B**, Start) of the theoretical natural isotope distribution, and the fully enriched spectrum (**Fig 1B**, Finish), parameterized by the n_L_ and (isotope molar percent enrichment, MPE), defines the theoretical distribution expected at infinite labeling time. Thus, for each neutromer (M_x_, X ∈ [M_0_, M_1_, M_2_, M_3_]), the fraction of new molecules (fraction new; FN) can be quantified as described in **Equation 2** by comparing its empirically observed intensity (I_x_) to its theoretical minimum (I_min_) and maximum (I_max_) labeling intensities.

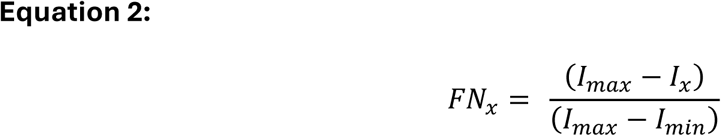

For each lipid and each time point, FN was calculated as the mean of per-neutromer FN values (FN_x_) across the most abundant neutromer peaks (**Equation 2**). This provided a mean FN and standard deviation for each molecule across the time series of *FN* values, *FN(t)*. The measurement standard deviation was used to filter for measurement quality as previously described ^13^, set to 0.1 for this study. Additionally, spectra with low signal intensity or minimal difference between fully labeled and unlabeled envelopes were excluded to ensure robust fits.

### Exponential-fit kinetic constants (k and A_syn_) from quality-filtered measurements

Final rate values (**Table S3 for peptides and S5 for lipids**) were obtained from filtered time-dependent FN(t) measurements for all related molecules fit in SciPy’s curve_fit function (v1.17) to an exponential replacement model (**Equation 3).** Agreement between the initial binomial-fit parameters and the final exponential-fit parameters was evaluated across high-quality lipid measurements, showing close correspondence for both turnover rate and A_syn_ estimates (**Fig S5**). For lipids, we observed that A_syn_ is highly variable as observed in previous literature ^28^, in contrast to the A_syn_ = 1 used for proteins ^15^.

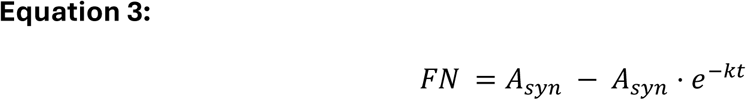

### Component *n_L_* derived from experimental lipid n_L_

Using the final *n_L_* for all detected proteins (**Fig 2**) or Lipids (**Fig 3**), we assembled a design matrix linking observed peptides or lipids to their molecular components. To ensure that only the highest-quality *n_L_* estimates were retained, we restricted our design matrix to peptides/lipids who’s empirical *n_L_* values did not exceed the total number of hydrogens (*n_H_*). We further included only those with covariance-derived standard errors < 3 for all three binomial fit parameters (B-k, B-A_syn_, and n_L_). Similar to the literature, for peptides A_syn_ in **Equation 1** was set to 1 ^13, 17, 20^ while for lipids we allowed A_syn_ to be variable ^28^.

**Figure 2:**
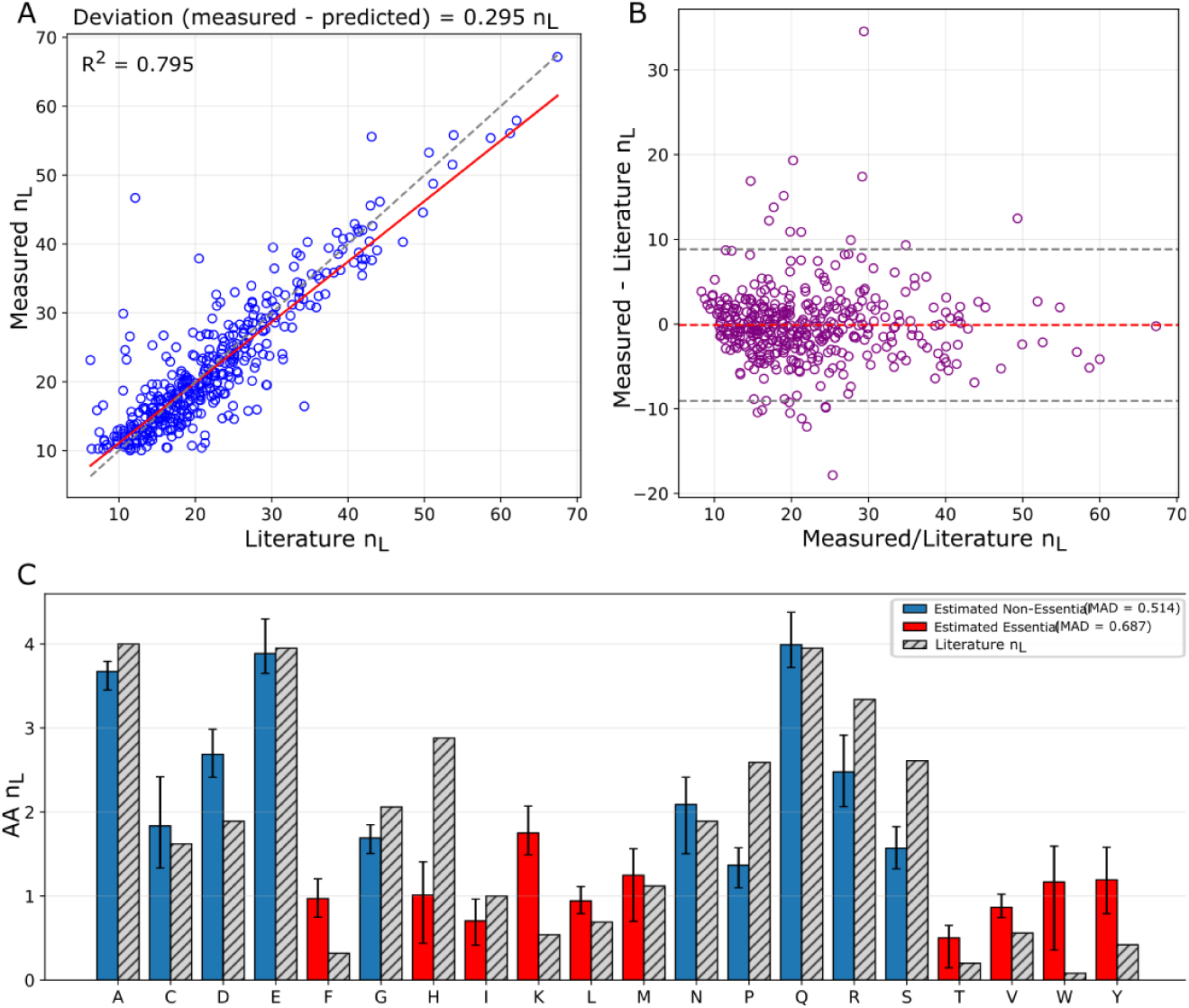
Peptide Positive Control. (A) Peptide-level measured literature n_L_ values trend (red) closely tracking the identity line (dashed). (B) Bland–Altman analysis of measured and predicted n_L_ values centered near zero across the dynamic range, indicating minimal systematic bias and acceptable dispersion. (C) Demixing of peptide to amino acid–level n_L_ values yields close agreement with theoretical expectations. Non-essential (blue) and essential (red) amino acids show low deviation from literature values (gray).

**Figure 3:**
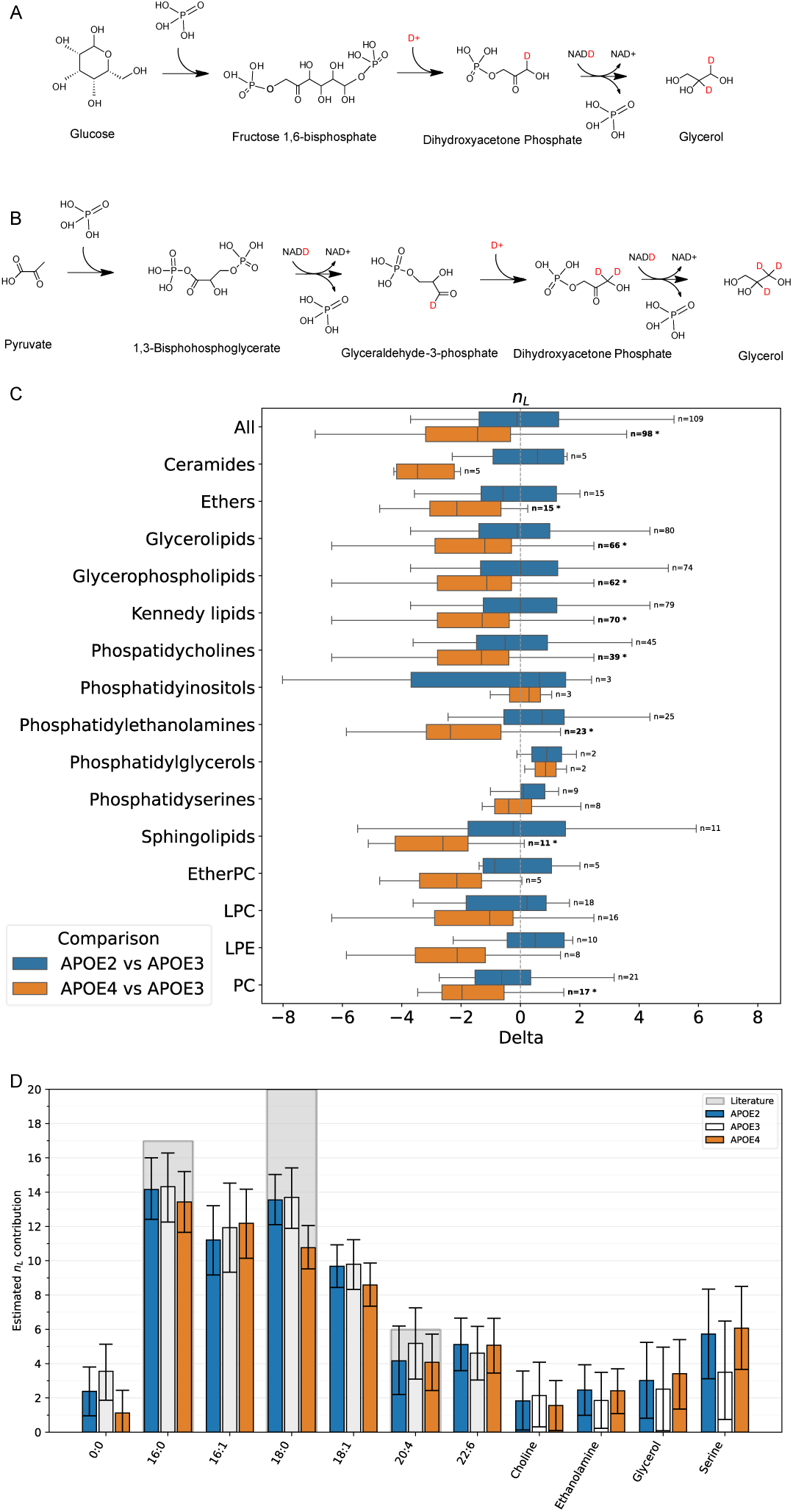
Non-integer n_L_ values reflect the combined contributions of multiple metabolic pathways: The literature n_L_ for glycerol is 2.5, representing contributions from dietary glucose with an n_L_ of 2 (A), and from dietary pyruvate with an n_L_ of 3 (B). Box plots of paired n_L_ differences for selected ontologies. An asterisk indicates corrected p < 0.05 (C). Estimated component n_L_ contributions for frequently observed lipid components with previously reported literature values in gray (D).

To propagate measurement uncertainty, we performed Monte Carlo demixing using standard errors derived from associated covariance matrices (**Fig S6**). We restricted the experimental lipid dataset to *n_L_* values below the theoretical hydrogen count and included only components detected in at least 3% of the lipids passing our quality cutoff filter. To assess analytical accuracy, we incorporated an artificial ‘0:0’ component into the design matrix for lyso and diacylglycerol species, which contain an unoccupied acyl chain position in place of a fatty acid (**Fig 3C**). This component carries no metabolically incorporated hydrogens and therefore has a definitionally true n_L_ of zero. The demixed 0:0 estimate therefore serves as a built-in accuracy diagnostic: because its true value is known to be zero, any deviation directly quantifies the magnitude of systematic error in the demixing procedure. We aggregated component-level estimates across Monte Carlo simulations and report the median n_L_ value with confidence intervals derived from the empirical distributions.

### Abundance Normalization

Two independent normalization methods for lipid abundance were used to investigate ApoE dependent-changes (**Fig S7**). Either an internal standard based or a median total ion chromatogram (mTIC) method were used. To enable standards-based normalization, we added a panel of deuterated (D7) lipid internal standards representing the major glycerophospholipid and neutral lipid classes at known final concentrations (phosphatidylcholine (PC), phosphatidylethanolamine (PE), phosphatidylglycerol (PG), Phosphatidylinositol (PI), Phosphatidylserine (PS), phosphatidic acid (PA), triacylglycerol (TG), sphingomyelin (SM), and cholesterol ester (CE); **Table S2**), from Avanti Lipids. The expected internal-standard species were consistently detected across samples, except for SM. For comparison of lipid quantities in the brain, we summed the DeuteRater-extracted neutromer intensities to minimize abundance bias caused by fraction-new-dependent decreases in M0 during deuterium incorporation. Specifically, M0–M2 were summed for lipid ions with m/z < 1400, while M0–M3 were summed for ions with m/z ≥ 1400, where the higher-mass isotope envelope required an additional neutromer to capture comparable unlabeled/labeled signal structure (**Fig. S10**).

For each data acquisition polarity, internal-standard intensities from every run were mapped to ApoE3 using a linear regression of the form **log_2_(y) = a + b·log_2_(x)**, where x denotes the ApoE3 intensity and y, the corresponding value from the ApoE2 or ApoE4 run. A Huber-loss function was used to limit outlier influence. The resulting parameters were applied to all lipid features, followed by back-transformation to linear space. For mTIC relative normalization all intensity values were log₂ transformed and the distribution of values for each sample was median centered to stabilize variance and linearize multiplicative technical effects ^38^. The mTIC approach scales summed neutromer intensities of each MS trace to an equivalent total ion signal, allowing proportional differences to be evaluated independently of total lipid content. Lipid ontologies were compared to ApoE3, following normalization using a one-sample hypothesis test on lipid-wise log₂ fold-changes corrected for multiple testing. For each lipid feature, the log₂ fold-change was computed as the difference between the ApoE2 or ApoE4 log₂-abundance and the corresponding ApoE3 value, yielding a log₂FC distribution for each genotype comparison. A two-sided one-sample t-test was then applied to determine whether the median log₂FC differed significantly from zero (H₀: median log₂FC = 0), using only features with finite, non-missing values. This was done for the same ontological groups as the kinetic variable paired t-tests.

### Kinetic variable comparisons

To evaluate genotype-dependent differences in kinetic variables (*k*, *A_syn_*, *n_L_*), we applied paired-sample hypothesis tests across several ontological groupings, including individual lipid classes (e.g., phosphatidylcholines (PC), phosphatidylethanolamines (PE), triacylglycerols (TG), cholesterol esters (CE), etc.), as well as higher-order categories such as Phospholipids, Glycerophospholipids, and Kennedy pathway (CDP-choline and CDP-ethanolamine) intermediates. For every lipid, a two-sided paired t-test was then applied to assess whether the median difference significantly deviated from zero (H₀: median difference = 0). For each kinetic variable, ApoE2 or ApoE4 values were paired with corresponding ApoE3 values for the same lipid feature. For these comparisons, the exponential decay fit R^2^ was greater than 0.6, the A_syn_ was between 0 and 1.2 in both datasets, and the *k* was within the range of 0.001 to 1 excluding lipids that were poorly constrained by the sampling times. For n_L_ comparisons we required that each had a value between 2 and the total number of hydrogens (n_H_) specified in the molecular formula.

To interpret coordinated changes in lipid abundance and turnover, we adapted the proteostasis framework described by Zuniga et al., 2024^15^ into a molecule-agnostic homeostasis analysis for metabolites. We assume that significant metabolite differences between an experimental condition and the control arise from metabolic regulation, in which changes in synthesis and/or degradation shift the system to a new steady state. In this model, both the initial and perturbed conditions are treated as locally stable states. Accordingly, a sufficiently large metabolic perturbation can move the system from one locally stable state to another, rather than returning to the original state through compensatory balancing of synthesis and degradation (**Fig 1A**).

The nature of this perturbation is inferred from the position of each metabolite in homeostasis space, where **log₂(relative fractional turnover)** is plotted against **log₂(relative abundance).** Lipids in quadrants I and III primarily reflect relative increases and decreases in synthesis, respectively, whereas lipids in quadrants II and IV indicate increases or decreases in degradation (Fig 5A).

Fractional turnover (k) is canonically defined as enzymatic flux divided by steady-state abundance (A), or **k = Flux/A**. Therefore, relative flux can be expressed as the product of relative turnover and relative abundance, where E is the experimental condition and C the control (**Equation 4**). Taking the log₂ transformation yields the additive relationship shown in **Equation 5**.

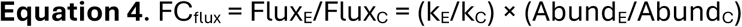

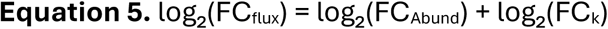

Given this relationship, the position of each metabolite relative to the line y = −x indicates constant flux. Points above the line correspond to increased inferred flux, interpreted as increased apparent pathway activity, whereas points below the line correspond to reduced inferred flux and lower apparent pathway activity. Statistical support for flux differences was assessed by combining abundance- and turnover-based p-values using Fisher’s combined chi-square test, yielding a χ ^2^ test statistic and a unified flux-associated p-value. When contextualized within the homeostasis plot, this approach enables detection of pathway activity changes while also indicating whether these shifts are more consistent with altered synthesis or altered degradation.

As a complementary analysis, we introduce the metabolic supply plot, in which **log₂(relative A_syn_)** is plotted against **log₂(relative abundance).** Here, A_syn_ represents the fraction of the lipid pool accessible to endogenous synthesis. Because the endogenously synthesized pool of each molecule is proportional to **abundance × A_syn_**, relative changes in endogenous synthesis can be expressed analogously to relative flux (**Equation 6**). Accordingly, the y = −x diagonal represents unchanged total endogenous synthesis between conditions. Points above this line indicate increased endogenous synthesis, whereas points below this line indicate reduced endogenous synthesis.

Metabolite placement within metabolite supply plot reveals whether changes in abundance are accompanied by an apparent synthetic response. In particular, analytes in quadrant IV are interpreted as reduced transport: these lipids are reduced in abundance and show no compensatory increase in percent-synthesis. Thus, continued depletion of these lipids may occur if abundance remains low without a corresponding increase in endogenous synthesis. Combined abundance–A_syn_ significance was assessed using Fisher’s combined chi-square test, analogous to the flux-based test used in homeostasis analysis.

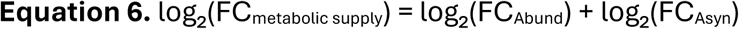

### Quantifying Covariance between kinetic parameters

For every lipid, the binomial fit (**Equation 1**) provided a covariance matrix (containing variances and covariances for n_L_, k, and A_syn_). From each matrix, we extracted and normalized the covariance between variables (**Fig S6**). For example, for n_L_ versus A_syn_ we normalized against the variance of Asyn to get the slope of the best-fit linear relationship between the two parameters as shown in **Equation 7** with A_syn_ as Variable1 and n_L_ as Variable2:

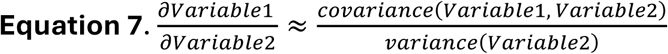

## Results

### Retention Time Matching

The MS1 scanning rate is an important part of MS1 spectral accuracy^13^, so for kinetics data acquisition no MS2 fragmentation was used to maximize MS1 acquisition. Implementation of our Legendre-based retention-time matching between MS2-identification runs and MS1-kinetics runs improved annotation of isotope patterns to lipid identifiers across all mass spectrometry data files (**Fig S1**). For example, two confidently identified positional isomers of the lysophosphatidylcholine 16:0 were identified. The extracted ion chromatogram (EIC) for this m/z was complex with multiple peaks within the correct retention time window. Further the RT shift between data acquisition runs was sufficient that there was an incorrect alignment of different isoforms at the MS1 level dramatically reducing data quality (**Fig S1A**). Using the unique signature of the EIC to align each of the observed peaks allowed us to generate sample specific retention times for each lipid. Applying these corrections improved isotopic-envelope fidelity (**Fig S4**), increasing the number of lipids that passed the DeuteRater-v7 quality-control filters. When a single reference retention time was used, 690 lipids between the three conditions had an exponential fit R^2^ > 0.6.

After the implementation of MS-file-specific retention times for each lipid, 731 had an R^2^ > 0.6. Although the aggregate improvement in relative deviation (RD) metrics is modest, the importance of the polynomial correction is most clearly illustrated in cases like lysophosphatidylcholine (LPC) 16:0, where two sn-positional isomers are only resolved after file-specific RT alignment, preventing incorrect co-extraction of isobaric species (**Fig S1**).

### Metabolic Isotope incorporation in peptides: *n_L_* calculation and demixing

As a proof of concept for the *n_L_* fitting and component *n_L_* demixing, we used a publicly available ^2^H_2_O-labeled peptide data set^13^ containing 24,822 measurements of 1,891 unique sequences from the liver of wild-type mice (**Table S3**). This data was collected using a QToF mass spectrometer which has higher spectral accuracy than other instruments.^13^ The peptide data were taken through the kinetic lipidomics workflow (**Fig 1D**) to derive all the relevant kinetic parameters (**Table S3**). Experimentally measured peptide-level *n_L_* values closely matched the literature n*_L_* values for each peptide, yielding an average difference of 0.3 and an overall linear correlation of R² = 0.80 (**Fig 2A**). Longer peptides with large n_L_ values had the largest deviation from the literature suggesting a small compounding overestimate in the sum of amino acid *n_L_* values based on the literature. The Bland Altman plot (**Fig 2B**) was centered on zero indicating low bias for peptide values.

We used linear algebra to “demix” component amino acid n_L_ values from these peptide sequences and compared to n_L_ library values (**Fig 2C**). There was no trend in accuracy per–amino acid size or chemistry or between essential and non-essential with median absolute deviation of 0.514 n_L_ and 0.687 n_L_ respectively. Histidine had the largest relative error in this analysis. The histidine error was likely not due to poor matrix constraints as histidine was observed in 418 unique sequences with thousands of measurements within the peptide data (**Table S4**). The variance could be due to instrumental noise or an increased dietary supply of unlabeled histidine ^39^. When the asymptote for the kinetic MIDA fit (Equation 1, kM-A_syn_) was allowed to vary (mirroring our lipidomics application) the average difference for full-peptide n_L_ values increased to 1.7 n_L_, with a corresponding decrease in linear correlation to R² = 0.47 (**Fig S8**). After allowing A_syn_ to vary, component-level amino acid-level median absolute deviations were 0.56 n_L_ and 0.191n_L_ for essential and non-essential amino acids, respectively. (**Table S4).**

### Metabolic Isotope incorporation in lipids: *n_L_* calculation and demixing

The n_L_ may change in measurable ways as metabolic pathways shift in response to precursor availability or enzyme concentrations (**Fig 3A,B**). In global comparisons of all the quantified lipids, the change in *n*_L_ for ApoE2 relative to ApoE3 was indiscernible from zero, while ApoE4 had a significant negative bias indicating smaller *n*_L_ values (d = -1.689, t = - 8.113, p = 1.54E-12; where d is Cohen’s effect size, t is the t-statistic, p is the Benjamini–Hochberg-corrected p-value for highly interrelated groups (**Table S6)**, i.e. the BH correction was applied to each lipid class separately (**Table S7)**. The negative bias in ApoE4 *n*_L_ was significant across most major lipid classes, including glycerophospholipids, glycerolipids, Kennedy pathway lipids, phosphatidylcholines, phosphatidylethanolamines, ethers, ceramides, and sphingolipids (**Fig 3C**). This suggests that the reduction in *n*_L_ in ApoE4 reflects a broadly distributed shift. Among these, ether lipids showed one of the more pronounced n*_L_* reductions in ApoE4, which is particularly interpretable given that plasmalogens resist gastrointestinal digestion and can enter circulation intact, making reduced ether lipid n*_L_* a potential indicator of increased peripheral lipid contribution to the brain lipidome.

Palmitic (16:0), stearic (18:0), and arachidonic (20:4) acid are three of the few fatty acids with previously reported *n_L_* values^11, 28^ (**Fig 3D, gray bars**). The literature *n_L_* for 20:4 is within the confidence interval of these measured values. The 16:0 n_L_ values in this study are ∼80% of the literature value ^11^. The 18:0 had almost the same n_L_ value which is 70% of the expected value. Because of the extra fatty acyl synthetase addition, it was expected that 18:0 should have a measured n_L_ value approximately 2-3 higher than 16:0. The value measured for 16:1 relative to 16:0 and 18:1 versus 18:0, does follow an expected pattern with both *n_L_* values decreasing by 2 due to the double bond. The essential fatty acids 20:4, 22:4, and 22:6 have very small *n_L_* values which are expected for these dietary sourced components. Additionally, glycerol and most headgroups are found in the expected range. The demixed 0:0 estimate was higher than zero, directly quantifying a small positive systematic error in the demixing procedure for this component. Choline and ethanolamine, both derived from serine, are around the expected n_L_ for serine (2.6), but serine itself is outside of the range.

Constituent-level n_L_ values were reproducible and did not show significant genotype-dependent shifts. This was notable because intact ApoE4 lipids showed a broad negative bias in n_L_ values. The component-level demixing estimates mass-averaged n_L_ values across all lipid classes while assuming a single well-mixed pool for each molecular component (A_syn_ = 1). Therefore, if a substantial fraction of a component, such as a fatty acid, entered the lipidome from dietary or otherwise unlabeled sources, that contribution would not be resolved directly in this component-level analysis. Instead, it may appear in the intact-lipid data as convolution between A_syn_ and n_L_. This parameter coupling is addressed further in the section “Quantifying Convolution in Kinetic Parameters.”

### Lipid Abundance Analysis

Protein concentrations in brain homogenates were used to normalize tissue input volumes and the amount of internal standard added to each sample. We therefore first evaluated whether internal-standard normalization removed technical differences among samples. As expected, the D7 internal standards showed tight agreement after normalization, with median log₂ fold-changes indistinguishable from zero in both ApoE2 vs. ApoE3 and ApoE4 vs. ApoE3 comparisons (**Fig. S7A,B**). However, when the same standard-derived correction was applied to the full lipidome there was a broad positive shift, most prominently in ApoE2 relative to ApoE3 and to a lesser extent in ApoE4 relative to ApoE3 (**Fig. S9**). Because sample input was normalized by protein concentration, we interpret this shift as consistent with genotype-associated differences in the lipid-to-protein ratio rather than a residual technical normalization error. Accordingly, internal-standard normalization produced highly significant positive abundance effects across many lipid ontologies, particularly in ApoE2. To minimize the influence of this global scaling effect and emphasize lipid-class remodeling relative to the overall lipidome, subsequent abundance analyses used mTIC-normalized values.^38,41^

Although not significant as with the standards normalization the mTIC-normalized ApoE2 abundances also had a bias towards higher abundances. Individual ontologies like phosphatidylcholines were modestly elevated (d = 0.234, t = 2.917, p = 0.004), while phosphatidic acids were reduced (d = −0.826, t = −4.176, p = 0.025) (**Fig 4A, blue**). The significant ApoE2 elevation in phosphatidylcholines is consistent with enhanced membrane biosynthetic activity, as PC is the predominant structural phospholipid in neuronal membranes and a key product of the Kennedy pathway. This may reflect the broader neuroprotective metabolic state associated with the ApoE2 isoform.

**Figure 4:**
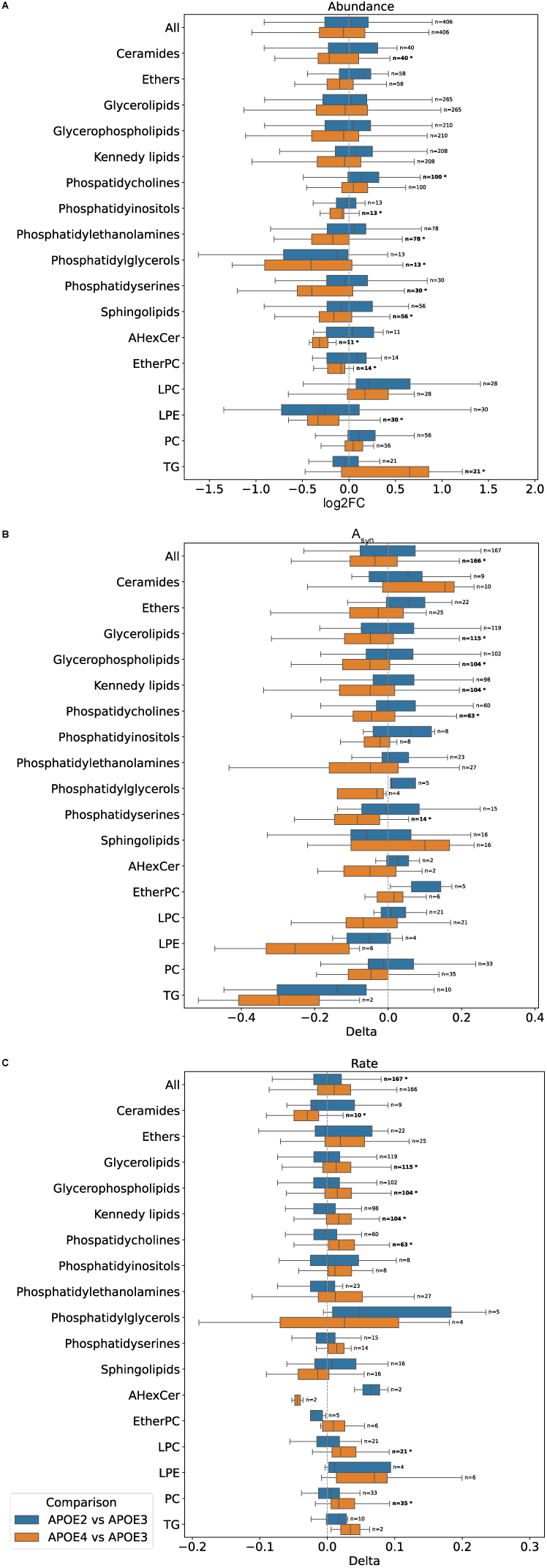
ApoE2 (blue) and ApoE4 (orange) versus ApoE3 across major ontologies. (A) Relative abundance (B) Biosynthesized fraction (A_syn_) shown as paired difference (Delta). (C) Fractional turnover rate (k) shown as paired difference (Delta). The number of paired comparisons per ontology is indicated by “n” to the right; bold n values with an asterisk indicate a Benjamini-corrected p < 0.05.

ApoE4 mTIC-normalized abundance revealed broader and more pronounced lipidomic changes than ApoE2 (**Fig. 4A, orange**). Sphingolipids (d = −0.167, t = −3.631, p = 0.001) and ceramides (d = −0.193, t = −3.506, p = 0.001) were significantly reduced, consistent with disrupted sphingolipid metabolism and potentially impaired endosome maturation, as identified in our previous proteomics study using the same mice.^15,46^ Several additional lipid groups, including phosphatidylethanolamines, phosphatidylserines, phosphatidylinositols, and phosphatidylglycerols, were also reduced (**Table S7**). The reduction in phosphatidylethanolamines (d = -0.186, t = -2.40, p = 0.0187) is notable given their enrichment in the inner mitochondrial membrane and the cytoplasmic leaflet of neuronal membranes, where their depletion may be the result of reduced mitochondria health. Intriguingly, acylcarnitines (CAR; d = −0.215, t = −75.415, p = 0.049, n = 2), which mediate fatty-acid trafficking into mitochondria, were also reduced, though the very small sample size warrants caution in interpreting this finding independently. In the context of increased triacylglycerols (TG; d = 0.499, t = 4.330, p < 0.001) and free fatty acids (FA; d = 0.853, t = 5.577, p = 0.005, n = 5), the co-occurrence of these observations is consistent with a model in which fatty acids are available but not efficiently imported into mitochondria for β-oxidation, instead accumulating as neutral lipid stores — though the low n-values for CAR and FA mean this mechanistic interpretation should be considered preliminary. Together, these observations are consistent with decreased mitochondrial health and with prior reports that ApoE4 promotes accumulation of neutral lipid species in the brain.^8,15,43–45^

### Biosynthesized Fraction: *A_syn_*

For proteomics experiments, there is consistently an *A_syn_* of 1 because of the fast degradation of dietary protein and nonessential amino acids ^13, 17, 20, 24, 25^. An *A_syn_* below one, indicates the presence of very slow-turnover biological sub-pools or significant dietary sourcing. These have been shown in previous studies of lipid metabolism ^5, 28, 40–43^. In ApoE2 tissue, the distribution of *A_syn_* values for all observed lipids was centered on zero (**Fig 4B**, blue) and no ontologies had significant shifts, suggesting that lipid population distributions are very similar to ApoE3.

Relative to ApoE3, we observed a small, but significant ApoE4-dependent decrease in A_syn_ across all lipids collectively (**Fig 4B, orange**, d = −0.041, t = −3.260, p = 0.001) and within multiple major lipid classes, including glycerophospholipids (d = −0.061, t = −4.353, p < 0.001), Kennedy pathway lipids (d = −0.065, t = −4.321, p < 0.001), glycerolipids (d = −0.059, t = −4.238, p < 0.001), phosphatidylserines (d = −0.093, t = −3.742, p = 0.002), and phosphatidylcholines (d = −0.057, t = −3.139, p = 0.003). We interpret the downward shift in A_syn_ primarily as the result of an increased proportion of the lipid pool arising from dietary contributions to fatty acids and other lipid constituents, rather than an increased influence from slow-turnover sub-pools. While accumulation of neutral lipid droplets has been reported in association with ApoE4 and may be linked to delayed endosomal maturation, this would require extensive synthesis of phospholipids using neutral lipids as precursors—a process that would increase rather than decrease A_syn_. Furthermore, degradation of neutral lipid stores would manifest as faster fractional turnover, which is observed, but without the corresponding reduction in abundance that a net depletion model would predict. Taken together, the ApoE4 A_syn_ phenotype is most consistent with an elevated dietary or peripheral lipid contribution to the measured pool.

### Turnover Rate: *k*

In ApoE4 tissue, we observed significant increases in the apparent turnover rate constant (*k*) across several lipid groupings (**Fig 4C, orange**), most prominently within Kennedy pathway lipids (mean difference = 0.0199, *t* = 3.43, *p* = 8.70 × 10^-4), and glycerophospholipids more broadly (mean difference = 0.0174, *t* = 3.28, *p* = 0.0014). This, taken in context with the co-reduced A_syn_ more generally, may support the notion of impaired blood-brain barrier in ApoE4.^54^ If full lipids were granted access to the central nervous system (CNS) from the blood, we might expect to see an increase in dietary sourcing, while simultaneously observing an increase in turnover rate as the slow-turnover lipids in the CNS co-mingle with the fast-turnover lipids in the blood. This ApoE4-dependent increase in turnover was not universal: ceramides exhibited a significant reduction in turnover rate (mean difference = -0.0305, *t* = -2.82, *p* = 0.0202), which can be interpreted as further support for the reduced endosome maturation phenotype.^15,46^

In ApoE2 tissue, the overall lipid pool showed a modest but significant increase in apparent turnover rate relative to ApoE3 (mean difference = 0.0210, *t* = 2.83, *p* = 0.0053). However, this shift was not concentrated within any individual ontology after multiple-testing correction. The strongest ontology-level effects, including non-hydroxy sphingosine ceramide (Cer_NS), dilysocardiolipin (DLCL), and ether triglycerides, did not remain significant after adjustment, indicating that the ApoE2-associated change in *k* is distributed across the lipidome rather than driven by a single lipid class.

### Quantifying convolution between kinetic parameters

The peptide control experiments highlighted that allowing *A*_syn_ to vary reduced the stability of amino acid-level n_L_ estimates (**Table S4**). Therefore, we quantified how the kinetic MIDA (kM) fit parameters co-varied in the binomial model (**Equation 1**). Each regression reports a Wald covariance matrix containing variances and covariances for n_L_, kM-*k*, and kM-*A*_syn_.

From each matrix, we extracted the covariance between each pair of parameters and normalized it by the variance of the second parameter, yielding a covariance-derived local sensitivity of Variable 1 with respect to Variable 2. For example, when kM-*k* is treated as Variable 1 and n_L_ as Variable 2, the resulting value estimates the expected change in kM-*k* per unit change in n_L_.

We projected these local sensitivities onto two-dimensional grids spanning the observed parameter space for each parameter pair (**Fig S6**). The resulting surfaces showed that parameter coupling was strongly directional. kM-*k* was minimally sensitive to n_L_ across the observed range (**Fig S6A**), whereas the reciprocal relationship showed elevated sensitivity of n_L_ to kM-*k* among slow-turnover species (**Fig S6B**). This indicates that, when turnover is poorly constrained, small uncertainty in rate can propagate into larger uncertainty in n_L_ even though n_L_ has little effect on the fitted rate.

A similar directional pattern was observed between kM-*k* and kM-*A*_syn_. kM-*k* showed only modest coupling to kM-*A*_syn_ across most of the observed parameter space (**Fig S6C**), whereas kM-*A*_syn_ became most sensitive to kM-*k* at very low turnover rates (**Fig S6D**). This rate-dependent coupling likely reflects the 32-day sampling window, which provides limited leverage to independently define both the turnover rate and asymptotic labeling plateau for extremely slow-turning-over species. Additional later timepoints would be expected to improve parameter identifiability in this regime.

The strongest evidence of parameter convolution was observed between kM-*A*_syn_ and n_L_. Negative coupling between kM-*A*_syn_ and n_L_ was most evident at higher kM-*A*_syn_ values and lower-to-intermediate n_L_ values (**Fig S6E**), indicating that increases in n_L_ are associated with lower fitted kM-*A*_syn_ in this region of parameter space. The reciprocal relationship was even more consequential: n_L_ was strongly and predominantly negatively sensitive to kM-*A*_syn_ across a broad intermediate-n_L_ band (**Fig S6F**). Thus, uncertainty in kM-*A*_syn_ can propagate substantially into n_L_ estimates. This analysis clarifies why allowing *A*_syn_ to vary expands confidence intervals for n_L_: the primary instability is not uniform coupling among all kinetic parameters, but directional propagation of uncertainty from rate and especially *A*_syn_ into n_L_ in specific regions of parameter space.

### Interpreting Lipid Homeostasis through Abundance and Turnover

Using the standards-based normalization for abundance removed the ontology nuances and made the flux and supply increases more extreme and global (**Fig S11**). Therefore homeostasis analyses used the mTIC-normalized abundance (**Fig 5 A, B**). The most impactful global observation for both ApoE2 and E4 was that small changes in biosynthetic fraction (Asyn, Fig 5B y-axis) or turnover rate (k, Fig 5A y-axis) accompany large differences in abundance (x-axis). The range of observed turnover rate changes was much smaller than for the protein turnover in these same samples.^15^

**Figure 5:**
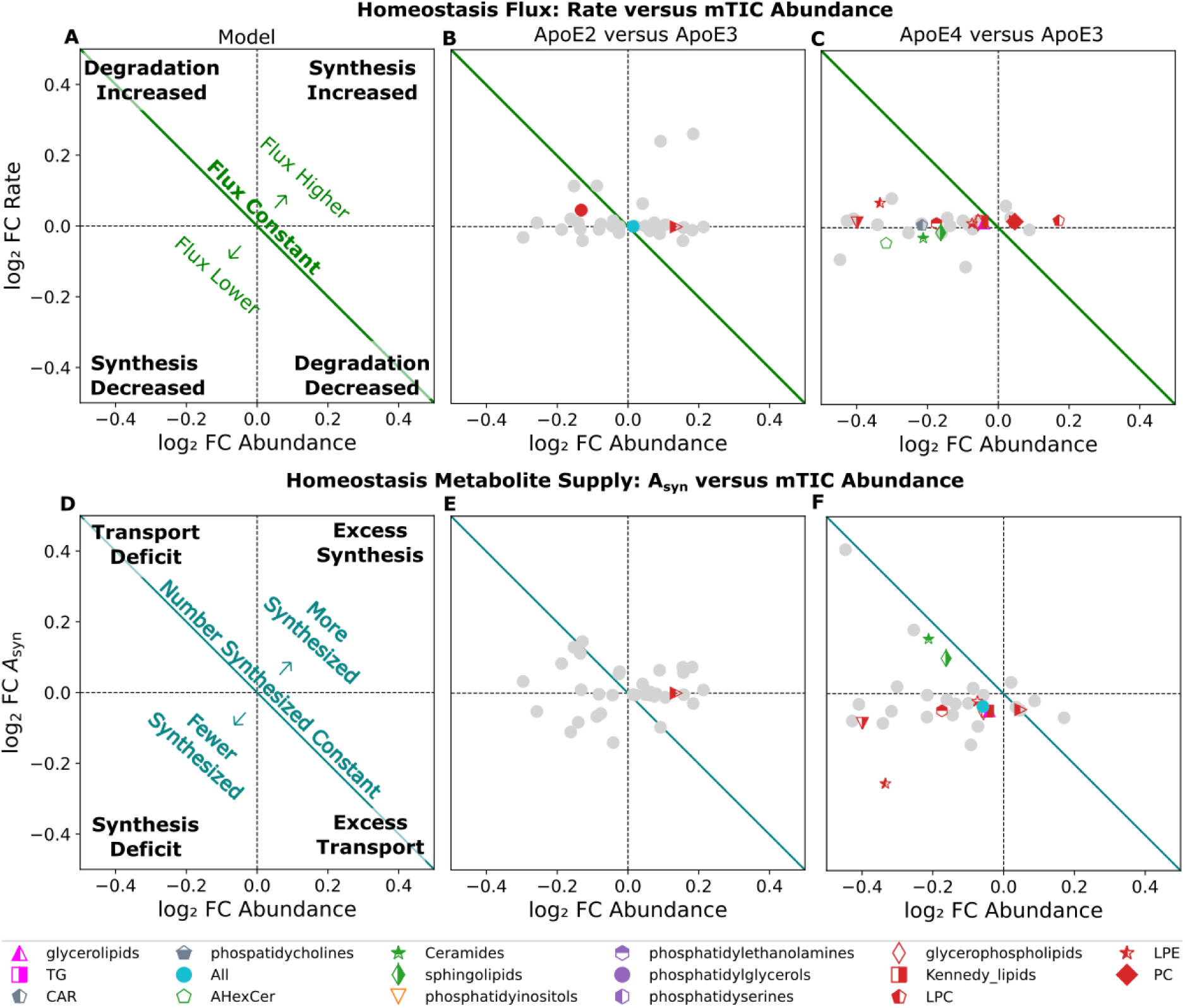
Flux (A–C) and metabolic supply (D–F) plots identify likely regulatory changes in lipid ontologies. Homeostasis plots show log_2_FC fractional turnover rate versus log_2_FC mTIC abundance; the green diagonal denotes equal flux and quadrants indicate changes in the relative ratio of synthesis to degradation. Metabolic supply plots show log_2_FC A_syn_ versus log_2_FC abundance; the diagonal denotes equal absolute synthesis, and quadrants the relative ratio of synthesis to transport. In both plot types, ontologies with significant changes in flux or total synthesis are colored; grey points represent ontologies that did not reach the Benjamini-Hochberg significance cutoff (p < 0.05).

Across all lipids, ApoE2 showed a small but significant increase in flux (the product of abundance and fractional turnover rate; median difference = 0.015, χ² = 13.432, p = 0.0094), due to a relative increase in synthesis. The strongest class-level effects were limited to increased flux in HighOrder phosphatidylcholines (median difference = 0.13, χ² = 11.076, p = 0.0257) with an apparent decrease in degradation. There was a reduced flux in HighOrder phosphatidylglycerols (median difference = −0.08, χ² = 9.709, adjusted p = 0.0456). Metabolite Supply analysis (**Fig 5 D, E**) similarly showed limited disruption, with HighOrder phosphatidylcholines being the only class with a significant positive combined shift (median difference = 0.14, χ² = 11.854, p = 0.0185) with an increase in transport.

Together, these results suggest that ApoE2 produces relatively constrained lipidomic remodeling, with the clearest response occurring in phosphatidylcholine metabolism.

In contrast, ApoE4 showed broader lipidomic disruption across both Homeostasis Flux and Metabolite Supply analyses (**Fig. 5C,F**), aligning with the organelle dysfunction observed in our companion proteomics study.¹⁵ There was a significantly reduced flux in HighOrder glycerophospholipids (median difference = −0.04, χ² = 18.749, p = 8.80 × 10⁻⁴) due to a relative decrease in degradation. Acylcarnitine (CAR median difference = −0.209, χ² = 13.075, p = 0.042, n = 2) also had significantly reduced flux due to an apparent increase in degradation. As discussed below, the interpretation of these changes may be convoluted by the low *n_L_*-value. Metabolite Supply analysis further identified passive reduction of glycerophospholipids (median difference = −0.107, χ² = 26.358, p = 2.68 × 10⁻⁵), indicating that these membrane-associated lipid pools are depleted and placing greater reliance on transport. HighOrder ceramides displayed significant changes across both analyses, with reduced flux (median difference = −0.24, χ² = 21.37, p = 2.27 × 10⁻⁴) driven by a reduced synthesis and passive diminishment (median difference = −0.121, χ² = 80.33, adjusted p = 0.0021), consistent with impaired late endosomal maturation.^15,46^ By contrast, triacylglycerols showed elevated flux and increased relative and total synthesis (Flux: median difference = 0.68, χ² = 17.690, p = 0.0109; Metabolite Supply: mean difference = 0.201, χ² = 17.868, p = 0.0150), suggesting that lipid substrates may be preferentially diverted toward neutral lipid storage rather than membrane repair or mitochondrial fatty-acid utilization. TG homeostasis and metabolic supply plot results could support prior evidence that human ApoE isoforms differentially affect brain glucose and ketone-body metabolism, as TG are primary involved in energy storage.⁴⁷ Together, these findings point toward a model in which ApoE4 promotes organelle vulnerability through aberrant delivery of lipid precursors and a failure to respond to low levels of important lipid molecules.

## Discussion

Unbiased shotgun assessments of lipid abundance have become a mainstay of metabolic research.^8,41,52^ Adding turnover rates to lipidome research dramatically expands metabolic insight^1,3,28,49^ by enabling quantification of the flux of molecules through enzymatic pathways.^1,3,28,53^ Many diseases incorporate modified lipid signaling and metabolism. Although mass spectrometry combined with ²H₂O labeling has been used to quantify turnover of individual lipids for decades,^10,29^ the complexity of isotope-distribution modeling^28^ has limited its extension to untargeted lipidomics.

As shown above, kinetic lipidomics measures turnover rate (*k*), endogenous synthesis fraction (*A*_syn_), and the number of isotope-incorporation sites (*n*_L_) for hundreds of identified lipids in a shotgun style experiment (**Fig 1C**). This provides a route to determine *n*_L_ in systems where reference libraries are unavailable, enabling high-throughput kinetic analysis. Because the approach does not require pre-existing *n*_L_ libraries, it should be adaptable to biological systems where ²H₂O labeling is feasible but isotope-incorporation behavior is poorly characterized. We demonstrated this method accurately measures well-documented^13,19,20,31^ amino acid *n*_L_ values in peptides (**Fig 2C**), enabling FN, *k*, and *A*_syn_ to be calculated in any experimental condition for both peptides and lipids.

After validating the fitting strategy, we used mice with genetically modified Apolipoprotein E (ApoE) as a biologically relevant perturbation to test whether kinetic lipidomics could resolve coordinated changes in lipid abundance, turnover, *A*_syn_, and *n*_L_. ApoE isoforms are of great interest because they can promote or protect against Alzheimer’s disease, although the mechanism is poorly understood. Multiple previous studies have investigated ApoE isoform effects on lipid abundance,^8,43,44,52^ but none examined turnover across the lipidome. The multi-parameter kinetic signature of ApoE4 brain tissue tells a coherent story that no single measurement could reveal alone (**Fig. 6**). We have previously shown with protein that the combination of abundance and turnover provides specific information about the relative contribution of synthesis to degradation in regulation of the proteome. Here we also show that the relative fraction of biosynthesis (*A_syn_*) can be combined with abundance to quantify the relative contribution of synthesis to transport across the lipidome.

**Figure 6:**
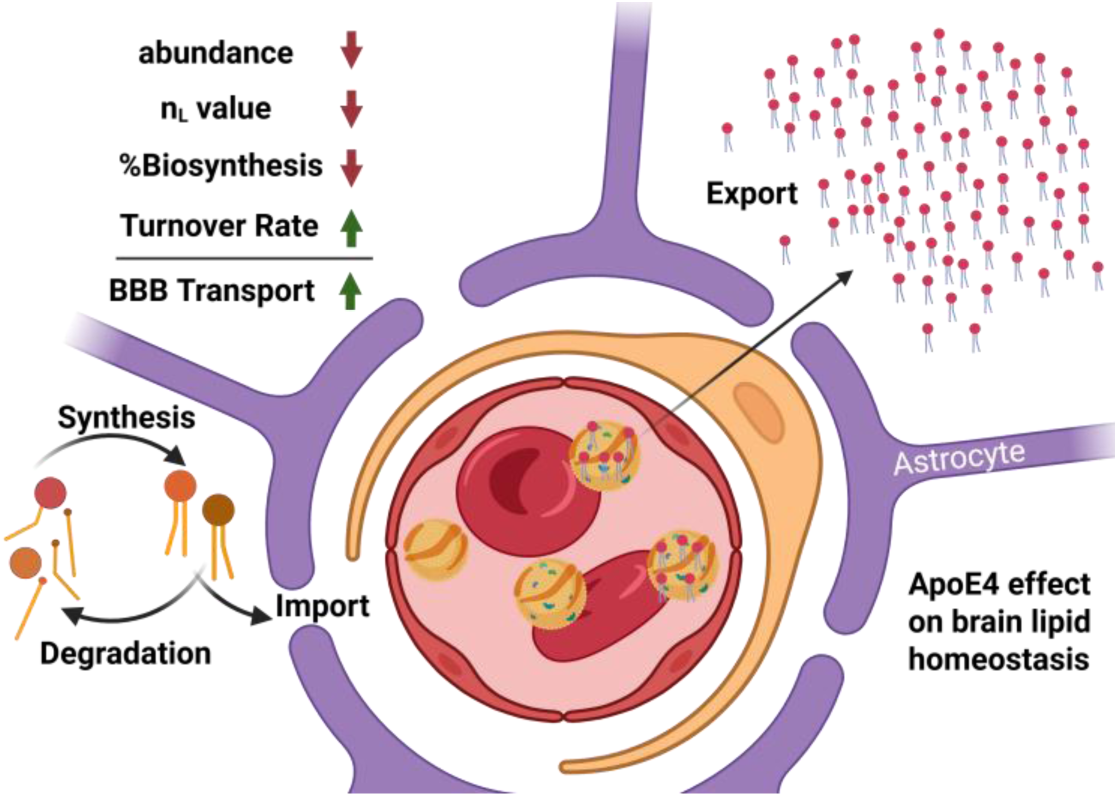
Model for metabolic perturbation caused by the ApoE isoform. Lipid metabolism is faster while incorporating more recycling and/or import of dietary sourced components consistent with a more permeable blood brain barrier in ApoE4

Our study included the protective isoform ApoE2. We found that both an internal-standard-based and an mTIC relative-distribution normalization of lipid abundance suggest that ApoE2 brain tissue has a meaningfully higher lipid-to-protein ratio than ApoE3, consistent with earlier literature reports^44–46^. Adding the kinetic analysis suggests that there is a greater overall flux in ApoE2 (**Fig 5, S11, Table S6**), which may underlie the neuroprotective effects of that isoform.

Here we show that the high-risk isoform ApoE4 exhibited broadly reduced lipid abundance, increased turnover rate, reduced *n*_L_, and lower *A*_syn_, particularly for lipid classes associated with mitochondria membranes and cellular maintenance, including phosphatidylethanolamines, ceramides, and acylcarnitines. Because dietary lipid components are unlabeled in this experimental design, reduced *n*_L_ across most major lipid ontologies (**Fig. 3C, Table S6**) is consistent with increased contribution from unlabeled stored or dietary pools. Reduced *A*_syn_ across glycerophospholipids, Kennedy pathway lipids, and phosphatidylcholines reinforces this interpretation: endogenous synthesis is contributing a smaller fraction of the measured pool. Reduced *n*_L_ in ether lipids is particularly informative in this context, as plasmalogens can resist gastrointestinal digestion and enter circulation intact,⁵⁵ making them a molecular indicator of peripheral lipid import into the brain. Together, the convergence of reduced *n*_L_ and reduced *A*_syn_ across most lipid classes supports the model shown in **Fig. 6**: ApoE4 brain tissue is receiving a larger unlabeled peripheral lipid contribution through a more permissive blood-brain barrier,⁵⁴ while simultaneously showing accelerated turnover that, for many membrane-associated lipid pools, is insufficient to compensate for ongoing losses.

This insufficiency is most clearly seen in specific lipid classes whose kinetic profiles reveal the underlying mechanisms. Ceramides showed reduced abundance, flux, and total endogenous synthesis (**Table S6**); ceramide supports intraluminal vesicle formation and exosome biogenesis within multivesicular endosomes⁴⁶ providing lipid-level evidence for the defective late endosomal trafficking reported by Zuniga et al.¹⁵ The concurrent reduction in CAR flux is consistent with impaired mitochondrial fatty-acid import and reduced β-oxidative capacity, which could lead to decreased mitochondrial health and the selective degradation of mitochondrial membrane proteins identified by Zuniga et al.^15^ Free fatty acids were also elevated in ApoE4 tissue (d = 0.853, t = 5.577, p = 0.049, n = 5), though the small sample size warrants caution. The co-occurrence of elevated free fatty acids and triacylglycerols alongside reduced CAR flux is consistent with a model in which fatty acids are available but not efficiently imported into mitochondria, instead being re-esterified into neutral lipid stores. By contrast, triacylglycerols showed elevated flux and total synthesis, seemingly the one lipid class moving in the opposite direction. This suggests that the ApoE4 lipidome is metabolically misrouted. Peripheral lipid import is elevated, membrane lipid synthesis is failing, and available fatty acids are diverted into neutral storage rather than supporting organelle maintenance.

## Conclusions and Future Work

This study establishes kinetic lipidomics as a practical, library-free method for measuring lipid turnover, endogenous synthesis fraction, and isotope-incorporation sites simultaneously across hundreds of lipid species. Applied to ApoE isoform mouse brain tissue, the method revealed that ApoE4 drives a broad, coordinated increase in turnover, reduced endogenous synthesis fraction, and reduced isotope-labeling sites across most major lipid classes. This seems to be consistent with increased neutral lipid storage and elevated peripheral lipid influx across a compromised blood–brain barrier. ApoE2 by contrast showed increased phosphatidylcholine flux with otherwise modest lipidomic remodeling relative to ApoE3. Validation against a published proteomics dataset confirmed that empirical n_L_ determination is accurate and generalizable, while the ApoE mouse cohort demonstrated that the method resolves biologically meaningful differences that static abundance measurements would miss entirely.

The ability to estimate n_L_ without reference libraries may be especially useful in systems where metabolic architecture differs substantially from mammalian models. Plant systems are a clear example, because lipid and amino acid biosynthetic pathways can diverge from mammalian labeling behavior and curated n_L_ libraries are generally unavailable. Because our framework can infer n_L_ without requiring a library, it is well suited for expanding kinetic lipidomics into plant biology and other non-mammalian systems where labeling behavior is poorly characterized.

A second priority is improving structural resolution within lipid classes, because unresolved fatty-acid isomers can obscure constituent-specific n_L_ assignments. Technologies such as aziridination, in-source ozonolysis, and epoxidation-based charge-inversion ion/ion chemistry could resolve double-bond positions and acyl-chain identities, enabling n_L_ values to be assigned at the level of individual fatty acyl chains. This would enable more precise n_L_ demixing for fatty acids and provide a framework for assessing how double-bond position and saturation state interact with lipid turnover and dietary sourcing.

## Limitations of this work

In contrast to isotope tracing which allows establishment of precursor and product relationships, this kinetic data is not pathway specific. Following up a kinetic lipidomics study with isotope tracing using purified precursors for the specific pathways of interest would provide complementary information. This study also represents the average changes over the entire brain. The contribution of regional or cell specific regulation should be addressed in future experiments.

## Acknowledgements

We are grateful for the assistance of the BYU Live Animal Facility, and the Fritz B. Burns Biological Mass Spectrometry Facility at BYU. This work was made possible by a grant from the National Institutes of Health [R01AG066874] to JCP; and Brigham Young University Undergraduate Research Awards to CRQ, KLV, KJC, SSG, MDS, MFP, TCH, BKD, ZH, and PMS. The funders had no role in study design, data collection and analysis, decision to publish, or preparation of the manuscript

## Data and Code Availability

The mass spectrometry data are publicly available via the Metabolomics Workbench partner repository with the dataset identifier **DataTrackID7141** This includes the raw LC-MS files used for quantitative and kinetic analysis in this study, as well as the MS-DIAL workflow and outputs. The DeuteRater-v7 code and an executable are available at GitHub: https://github.com/JC-Price/DeuteRater The Kinetic Lipidomics workflow is available on GitHub https://github.com/JC-Price/DeuteRater_Tools/tree/main/Kinetic_Lipidomics

