## Supplemental Figures for "Kinetic Lipidomics: Quantifying *in vivo* changes in lipid metabolism using metabolic labeling"

### Slide 1
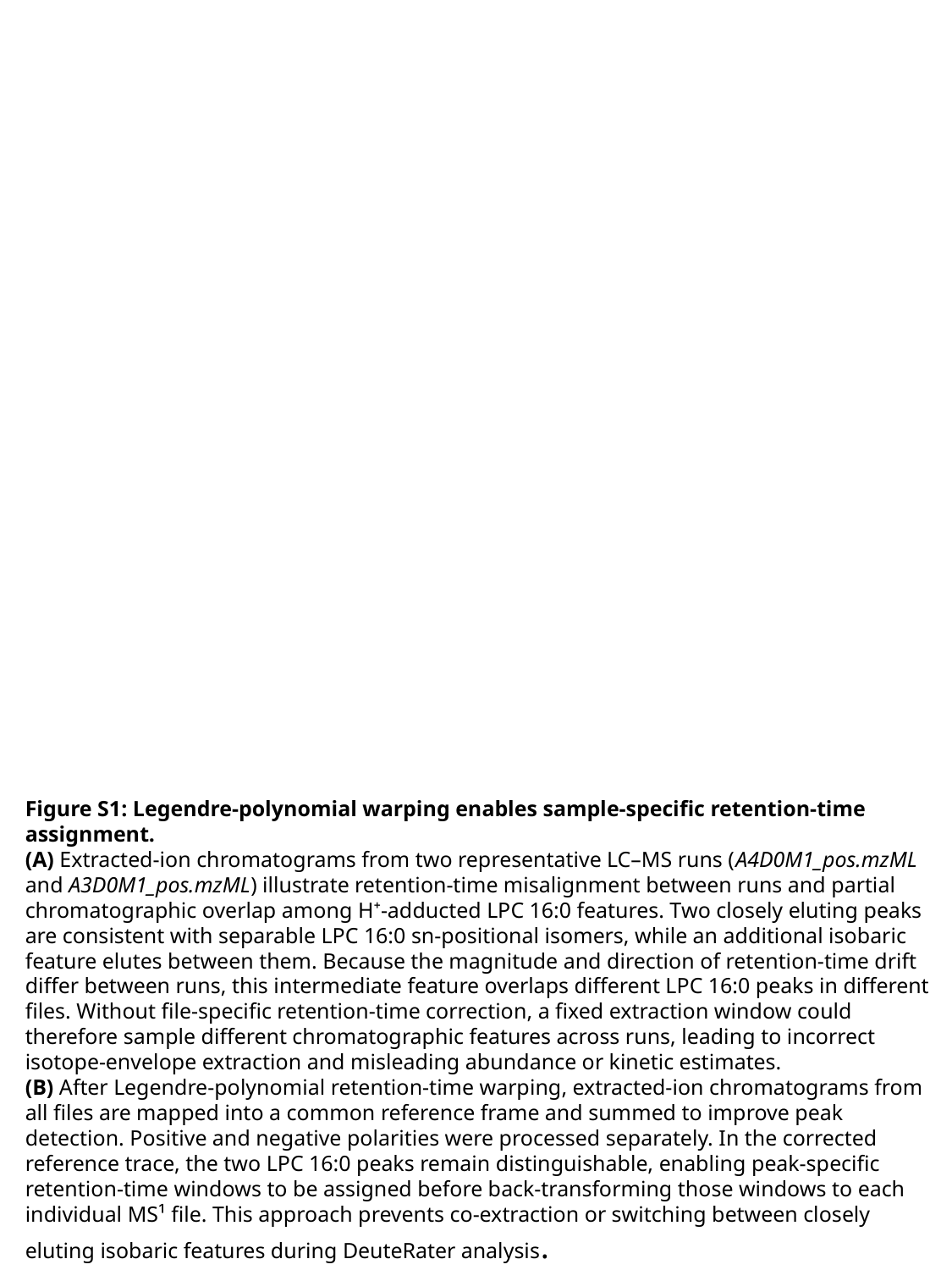

Figure S1: Legendre-polynomial warping enables sample-specific retention-time assignment.
(A) Extracted-ion chromatograms from two representative LC–MS runs (A4D0M1_pos.mzML and A3D0M1_pos.mzML) illustrate retention-time misalignment between runs and partial chromatographic overlap among H⁺-adducted LPC 16:0 features. Two closely eluting peaks are consistent with separable LPC 16:0 sn-positional isomers, while an additional isobaric feature elutes between them. Because the magnitude and direction of retention-time drift differ between runs, this intermediate feature overlaps different LPC 16:0 peaks in different files. Without file-specific retention-time correction, a fixed extraction window could therefore sample different chromatographic features across runs, leading to incorrect isotope-envelope extraction and misleading abundance or kinetic estimates.
(B) After Legendre-polynomial retention-time warping, extracted-ion chromatograms from all files are mapped into a common reference frame and summed to improve peak detection. Positive and negative polarities were processed separately. In the corrected reference trace, the two LPC 16:0 peaks remain distinguishable, enabling peak-specific retention-time windows to be assigned before back-transforming those windows to each individual MS¹ file. This approach prevents co-extraction or switching between closely eluting isobaric features during DeuteRater analysis.

### Slide 2
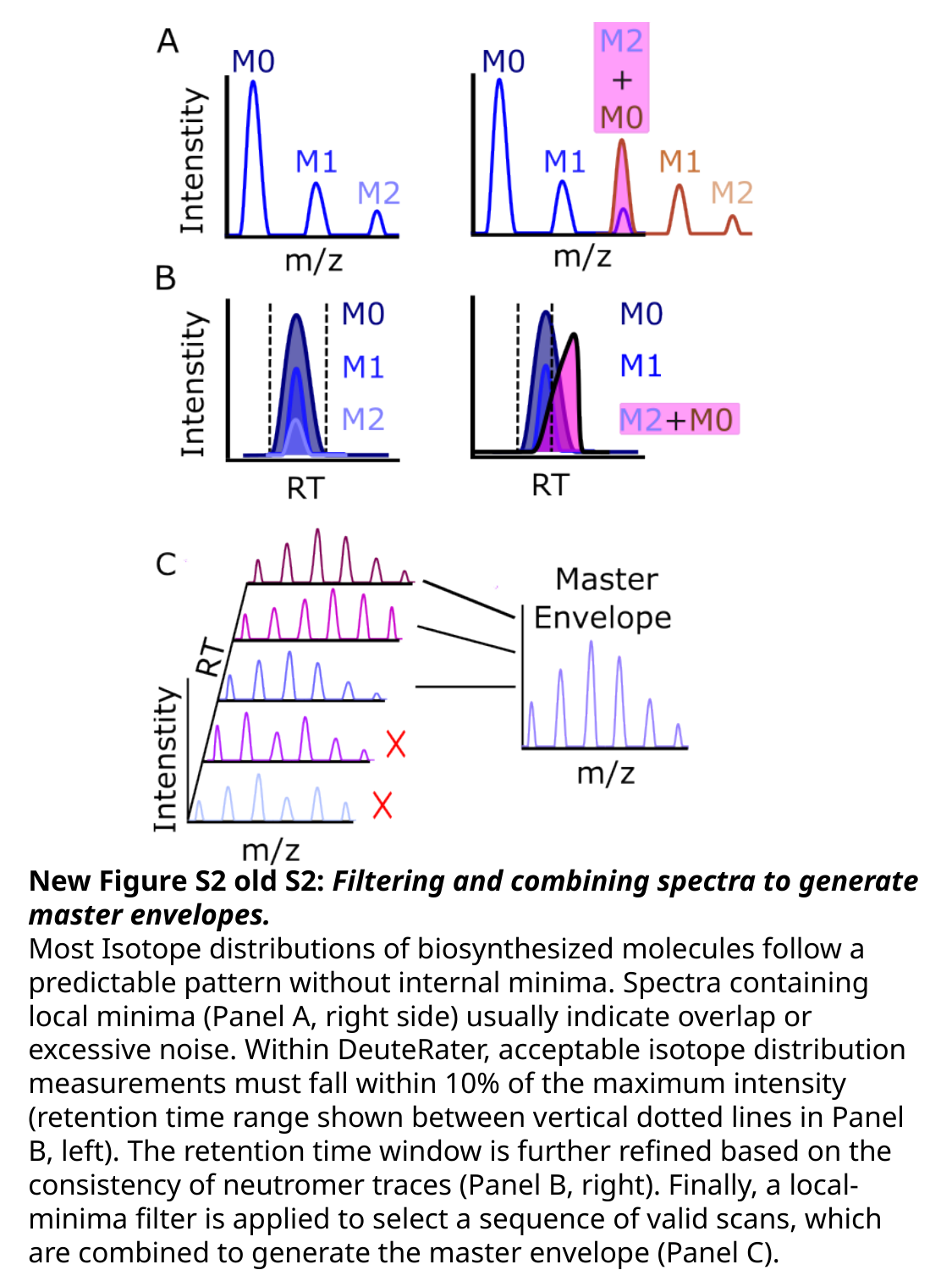

New Figure S2 old S2: Filtering and combining spectra to generate master envelopes.
Most Isotope distributions of biosynthesized molecules follow a predictable pattern without internal minima. Spectra containing local minima (Panel A, right side) usually indicate overlap or excessive noise. Within DeuteRater, acceptable isotope distribution measurements must fall within 10% of the maximum intensity (retention time range shown between vertical dotted lines in Panel B, left). The retention time window is further refined based on the consistency of neutromer traces (Panel B, right). Finally, a local-minima filter is applied to select a sequence of valid scans, which are combined to generate the master envelope (Panel C).

### Slide 3
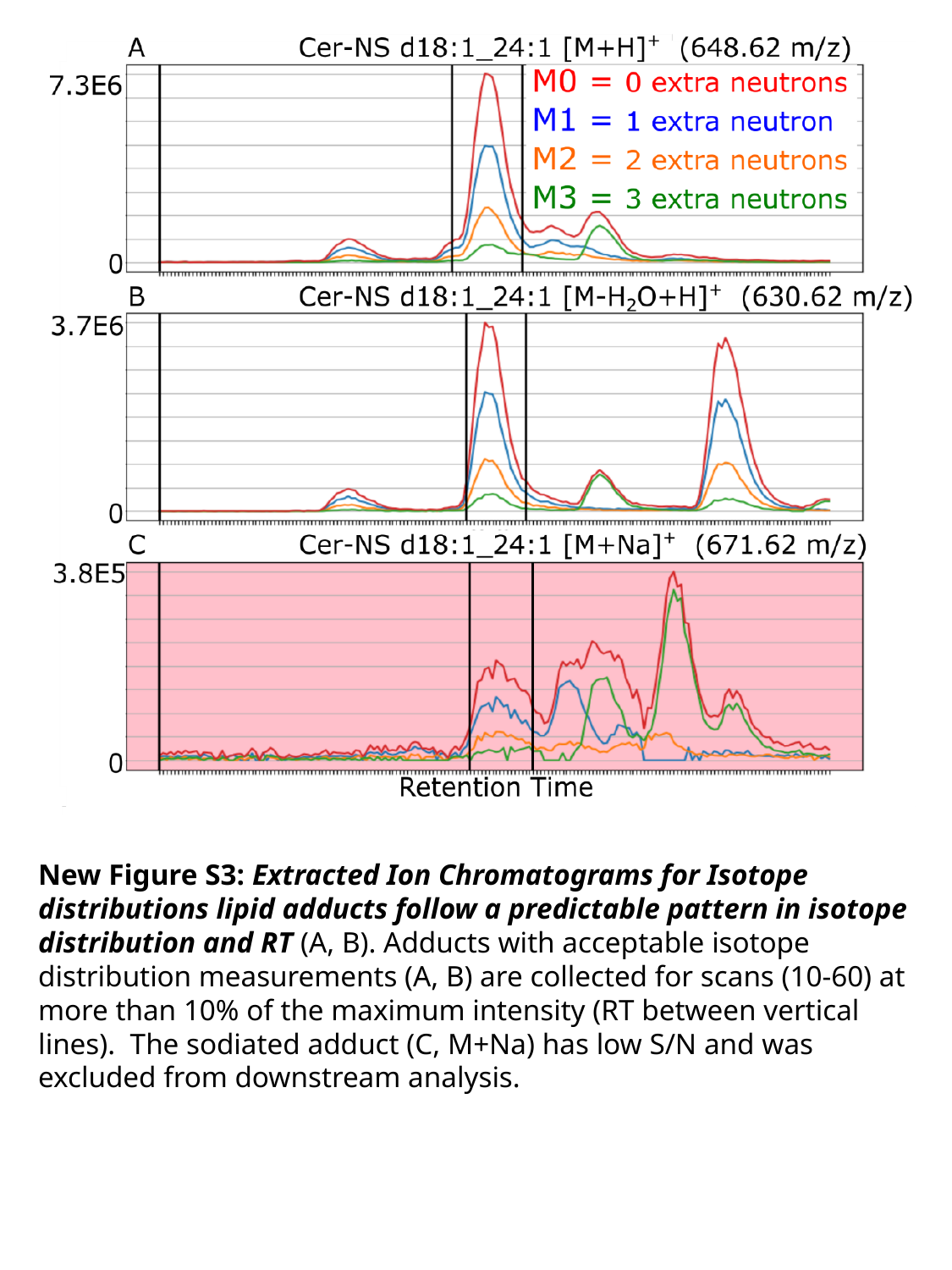

New Figure S3: Extracted Ion Chromatograms for Isotope distributions lipid adducts follow a predictable pattern in isotope distribution and RT (A, B). Adducts with acceptable isotope distribution measurements (A, B) are collected for scans (10-60) at more than 10% of the maximum intensity (RT between vertical lines). The sodiated adduct (C, M+Na) has low S/N and was excluded from downstream analysis.

### Slide 4
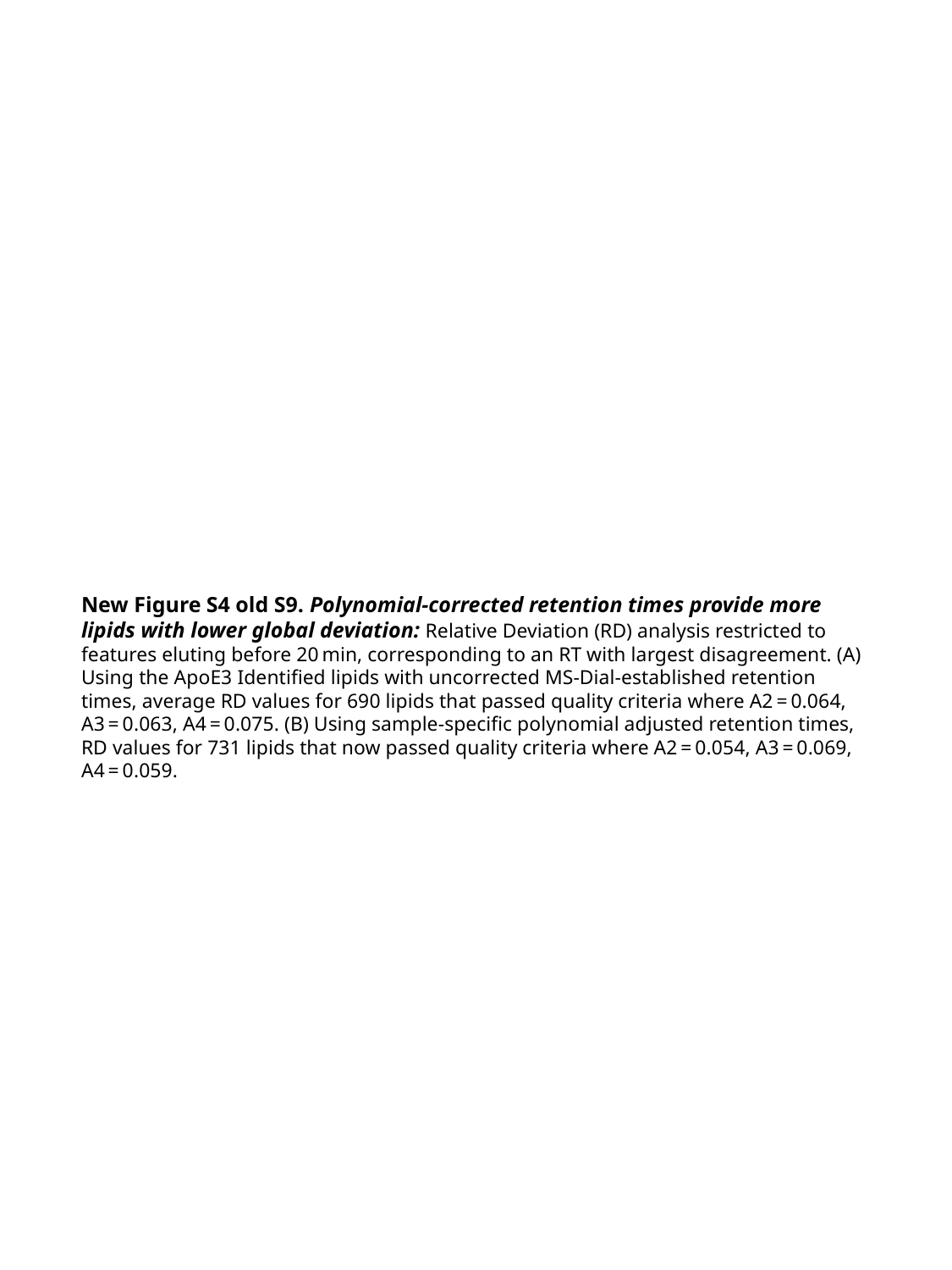

New Figure S4 old S9. Polynomial-corrected retention times provide more lipids with lower global deviation: Relative Deviation (RD) analysis restricted to features eluting before 20 min, corresponding to an RT with largest disagreement. (A) Using the ApoE3 Identified lipids with uncorrected MS-Dial-established retention times, average RD values for 690 lipids that passed quality criteria where A2 = 0.064, A3 = 0.063, A4 = 0.075. (B) Using sample-specific polynomial adjusted retention times, RD values for 731 lipids that now passed quality criteria where A2 = 0.054, A3 = 0.069, A4 = 0.059.

### Slide 5
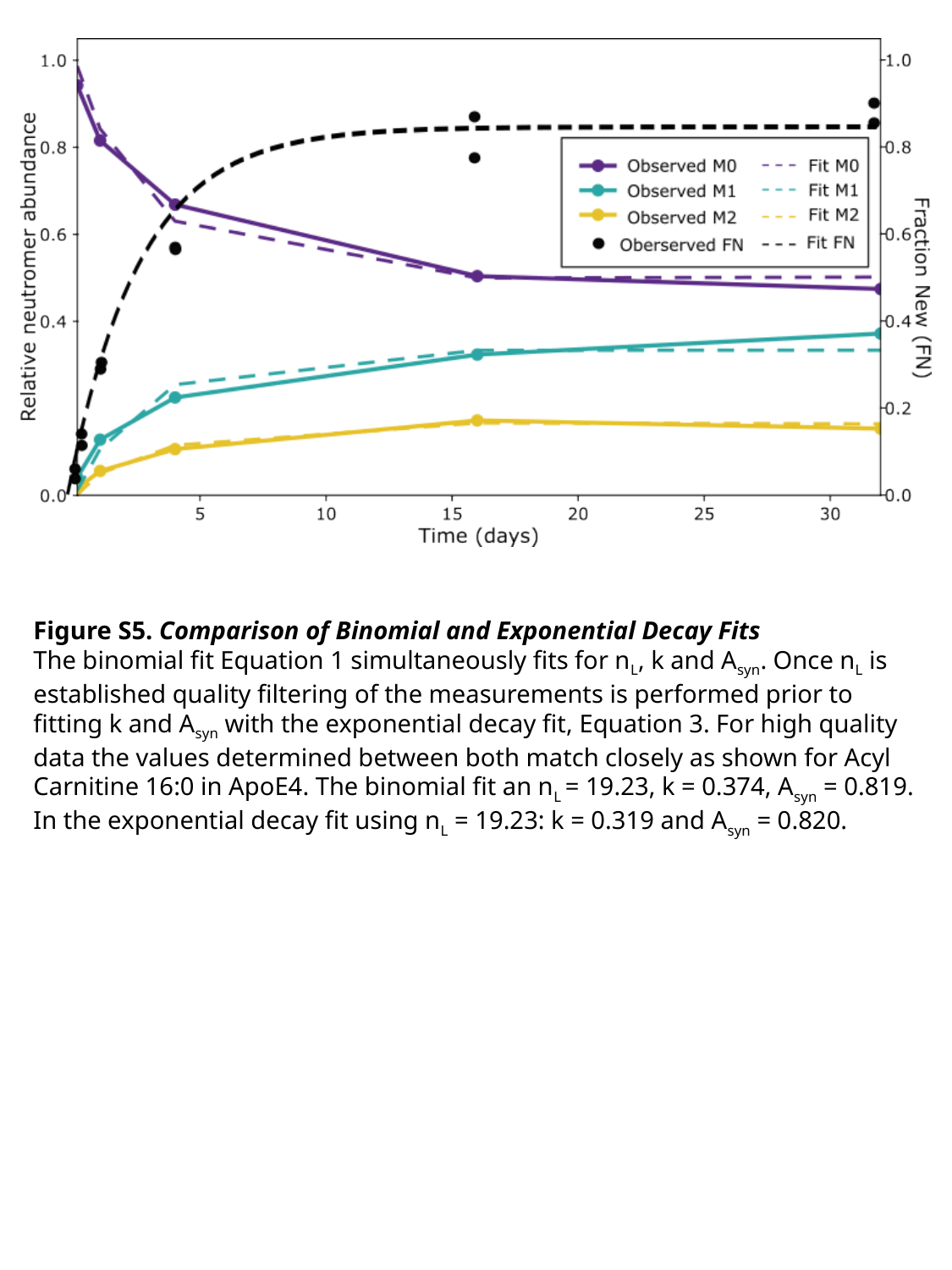

Figure S5. Comparison of Binomial and Exponential Decay Fits
The binomial fit Equation 1 simultaneously fits for n_L_, k and A_syn_. Once n_L_ is established quality filtering of the measurements is performed prior to fitting k and A_syn_ with the exponential decay fit, Equation 3. For high quality data the values determined between both match closely as shown for Acyl Carnitine 16:0 in ApoE4. The binomial fit an n_L_ = 19.23, k = 0.374, A_syn_ = 0.819. In the exponential decay fit using n_L_ = 19.23: k = 0.319 and A_syn_ = 0.820.

### Slide 6
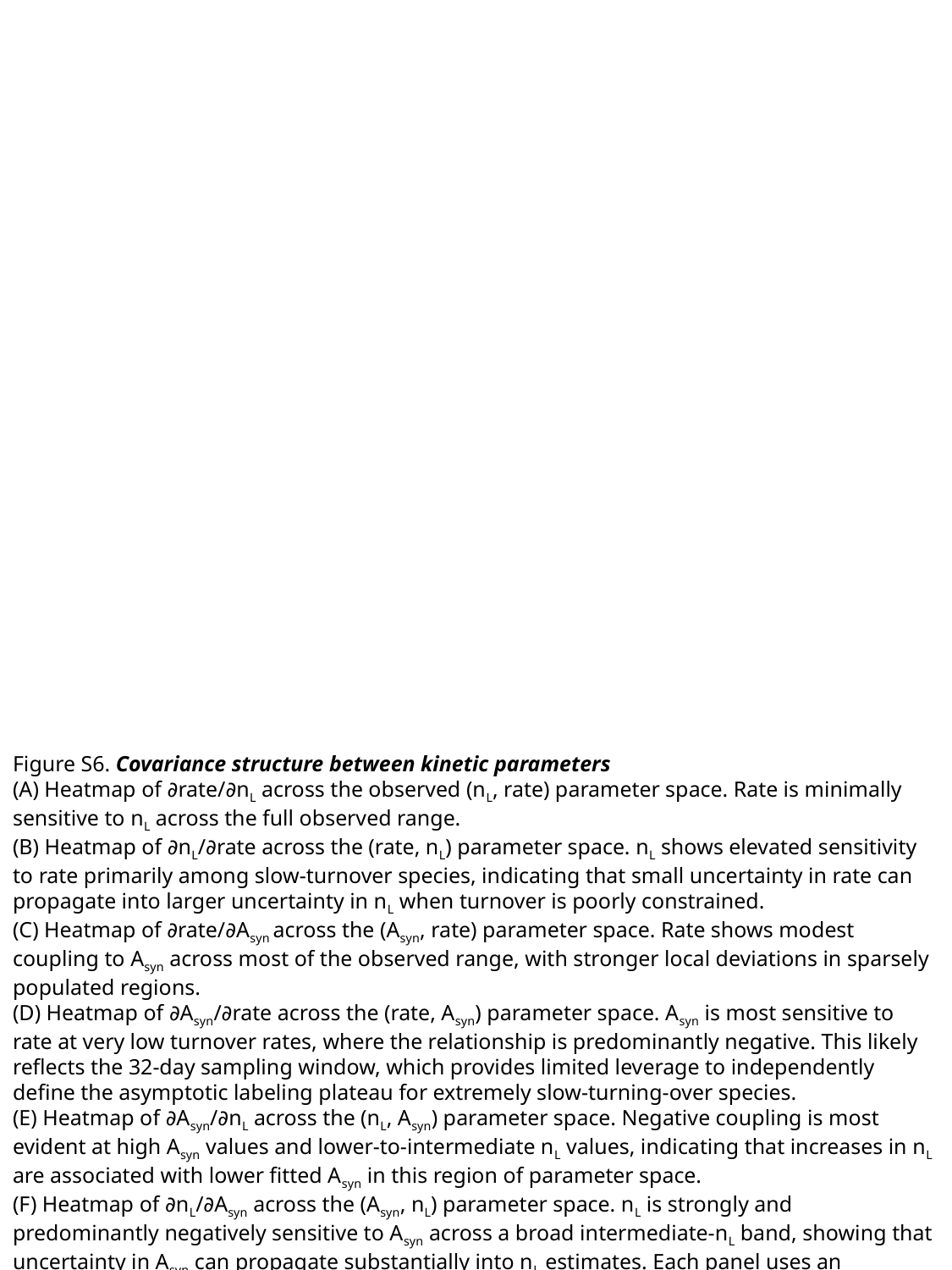

Figure S6. Covariance structure between kinetic parameters
(A) Heatmap of ∂rate/∂n_L_ across the observed (n_L_, rate) parameter space. Rate is minimally sensitive to n_L_ across the full observed range.
(B) Heatmap of ∂n_L_/∂rate across the (rate, n_L_) parameter space. n_L_ shows elevated sensitivity to rate primarily among slow-turnover species, indicating that small uncertainty in rate can propagate into larger uncertainty in n_L_ when turnover is poorly constrained.
(C) Heatmap of ∂rate/∂A_syn_ across the (A_syn_, rate) parameter space. Rate shows modest coupling to A_syn_ across most of the observed range, with stronger local deviations in sparsely populated regions.
(D) Heatmap of ∂A_syn_/∂rate across the (rate, A_syn_) parameter space. A_syn_ is most sensitive to rate at very low turnover rates, where the relationship is predominantly negative. This likely reflects the 32-day sampling window, which provides limited leverage to independently define the asymptotic labeling plateau for extremely slow-turning-over species.
(E) Heatmap of ∂A_syn_/∂n_L_ across the (n_L_, A_syn_) parameter space. Negative coupling is most evident at high A_syn_ values and lower-to-intermediate n_L_ values, indicating that increases in n_L_ are associated with lower fitted A_syn_ in this region of parameter space.
(F) Heatmap of ∂n_L_/∂A_syn_ across the (A_syn_, n_L_) parameter space. n_L_ is strongly and predominantly negatively sensitive to A_syn_ across a broad intermediate-n_L_ band, showing that uncertainty in A_syn_ can propagate substantially into n_L_ estimates. Each panel uses an independently auto-scaled symmetric log color scale centered at zero; color intensity should therefore be interpreted within each panel rather than compared directly across panels.

### Slide 7
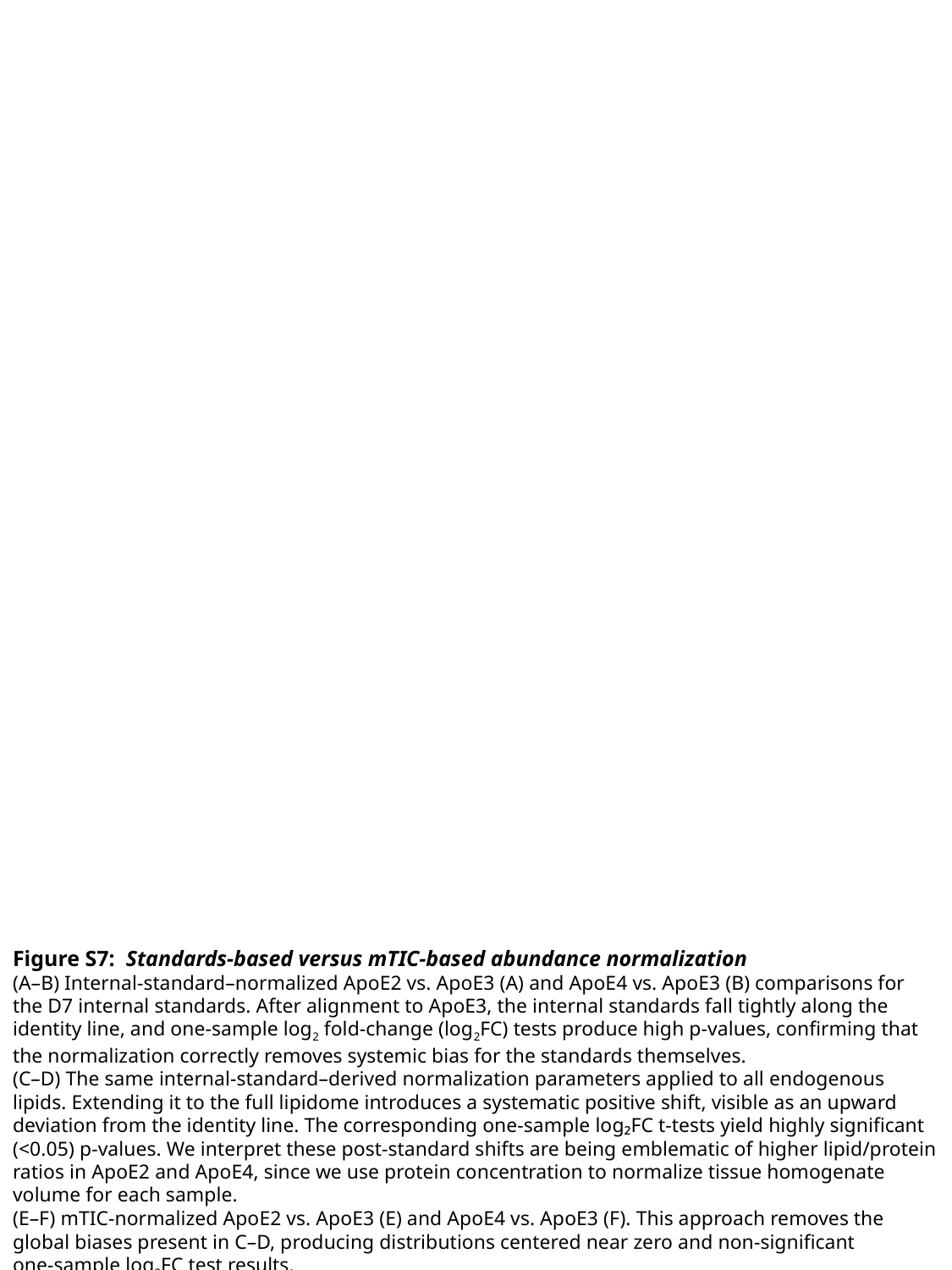

Figure S7: Standards-based versus mTIC-based abundance normalization
(A–B) Internal‑standard–normalized ApoE2 vs. ApoE3 (A) and ApoE4 vs. ApoE3 (B) comparisons for the D7 internal standards. After alignment to ApoE3, the internal standards fall tightly along the identity line, and one‑sample log_2_ fold‑change (log_2_FC) tests produce high p‑values, confirming that the normalization correctly removes systemic bias for the standards themselves.
(C–D) The same internal‑standard–derived normalization parameters applied to all endogenous lipids. Extending it to the full lipidome introduces a systematic positive shift, visible as an upward deviation from the identity line. The corresponding one‑sample log₂FC t‑tests yield highly significant (<0.05) p‑values. We interpret these post-standard shifts are being emblematic of higher lipid/protein ratios in ApoE2 and ApoE4, since we use protein concentration to normalize tissue homogenate volume for each sample.
(E–F) mTIC‑normalized ApoE2 vs. ApoE3 (E) and ApoE4 vs. ApoE3 (F). This approach removes the global biases present in C–D, producing distributions centered near zero and non‑significant one‑sample log_2_FC test results.

### Slide 8
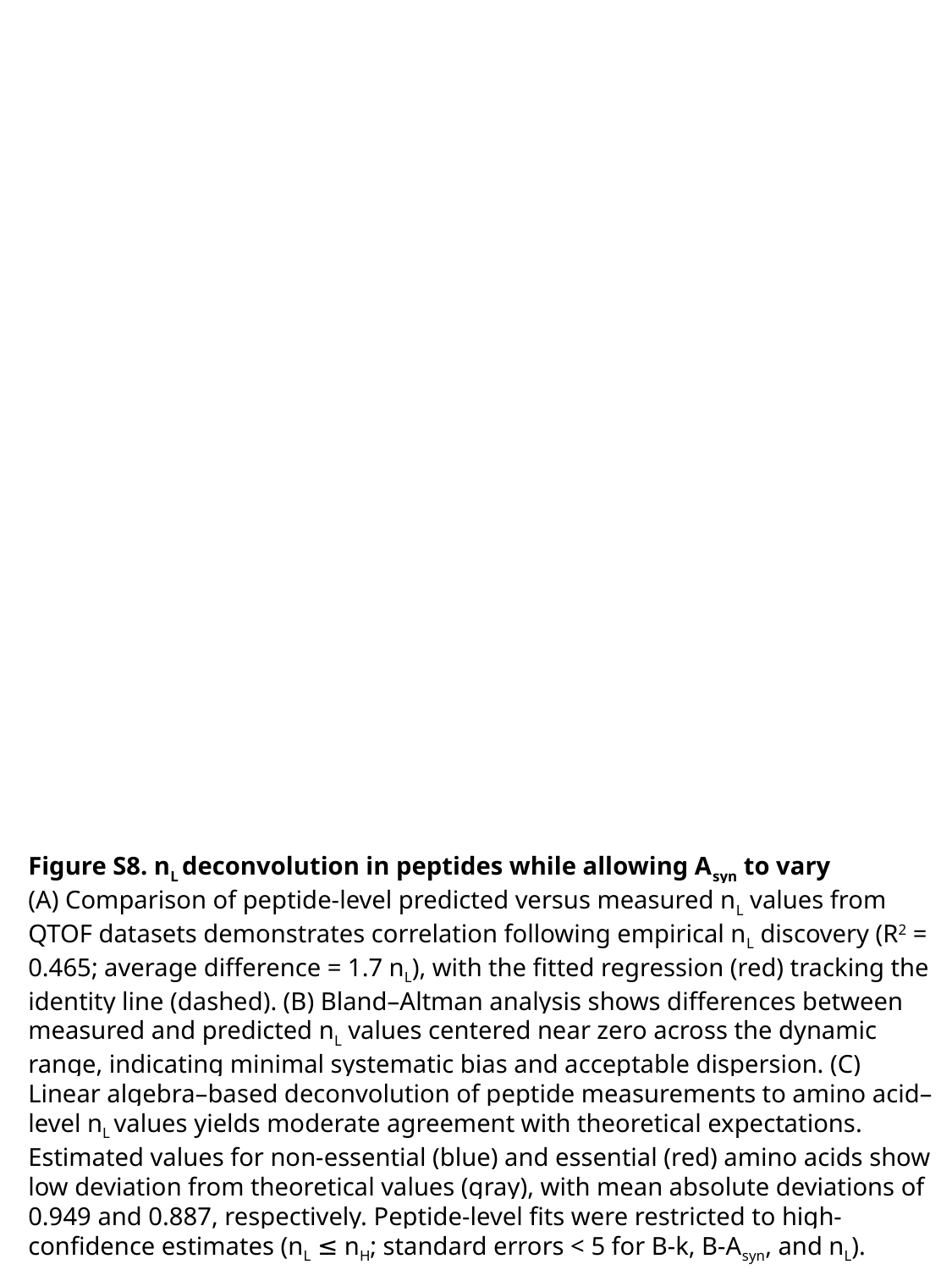

Figure S8. n_L_ deconvolution in peptides while allowing A_syn_ to vary
(A) Comparison of peptide-level predicted versus measured n_L_ values from QTOF datasets demonstrates correlation following empirical n_L_ discovery (R^2^ = 0.465; average difference = 1.7 n_L_), with the fitted regression (red) tracking the identity line (dashed). (B) Bland–Altman analysis shows differences between measured and predicted n_L_ values centered near zero across the dynamic range, indicating minimal systematic bias and acceptable dispersion. (C) Linear algebra–based deconvolution of peptide measurements to amino acid–level n_L_ values yields moderate agreement with theoretical expectations. Estimated values for non-essential (blue) and essential (red) amino acids show low deviation from theoretical values (gray), with mean absolute deviations of 0.949 and 0.887, respectively. Peptide-level fits were restricted to high-confidence estimates (n_L_ ≤ n_H_; standard errors < 5 for B-k, B-A_syn_, and n_L_).

### Slide 9
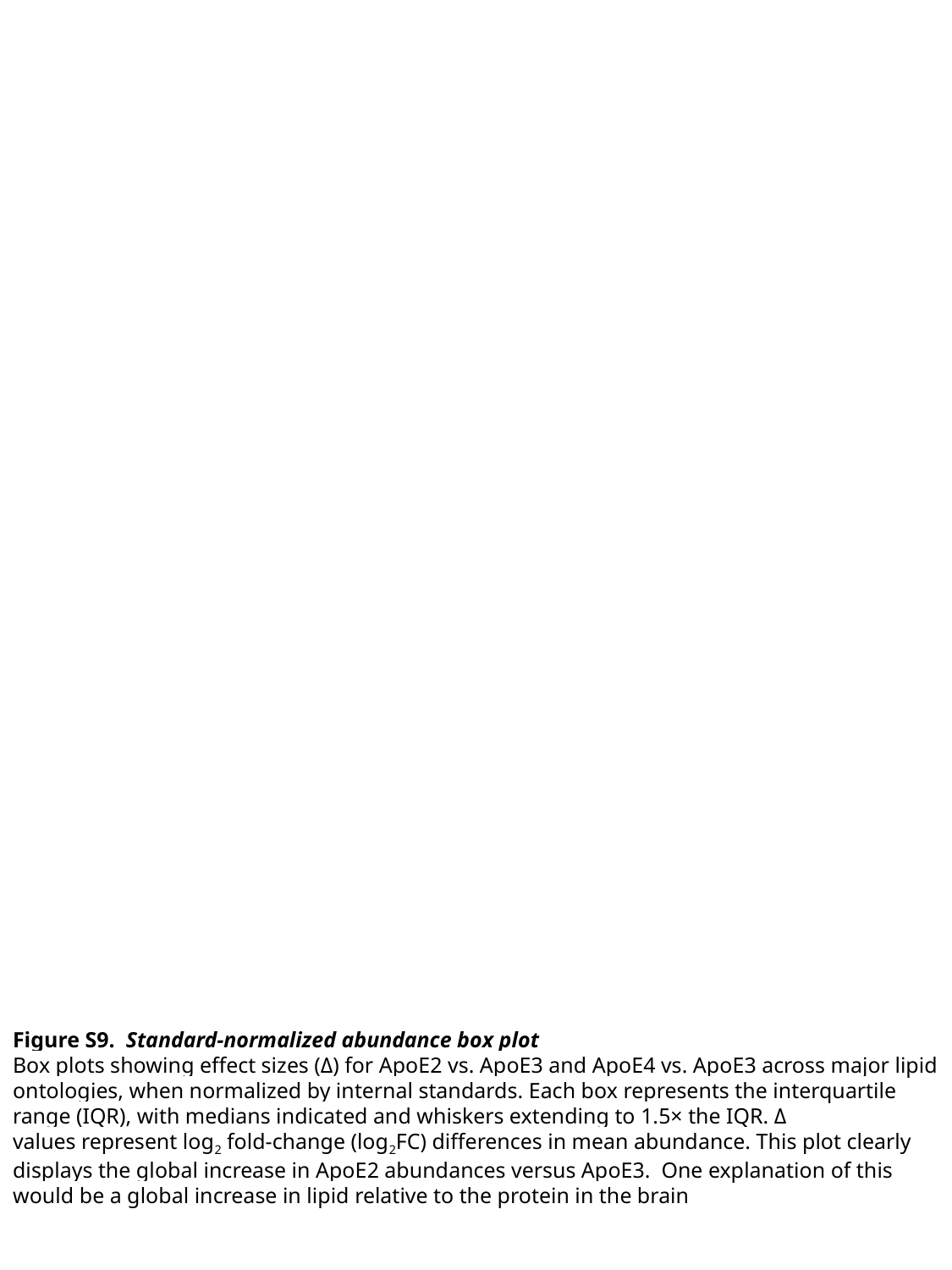

Figure S9. Standard-normalized abundance box plot
Box plots showing effect sizes (Δ) for ApoE2 vs. ApoE3 and ApoE4 vs. ApoE3 across major lipid ontologies, when normalized by internal standards. Each box represents the interquartile range (IQR), with medians indicated and whiskers extending to 1.5× the IQR. Δ values represent log_2_ fold‑change (log_2_FC) differences in mean abundance. This plot clearly displays the global increase in ApoE2 abundances versus ApoE3. One explanation of this would be a global increase in lipid relative to the protein in the brain

### Slide 10
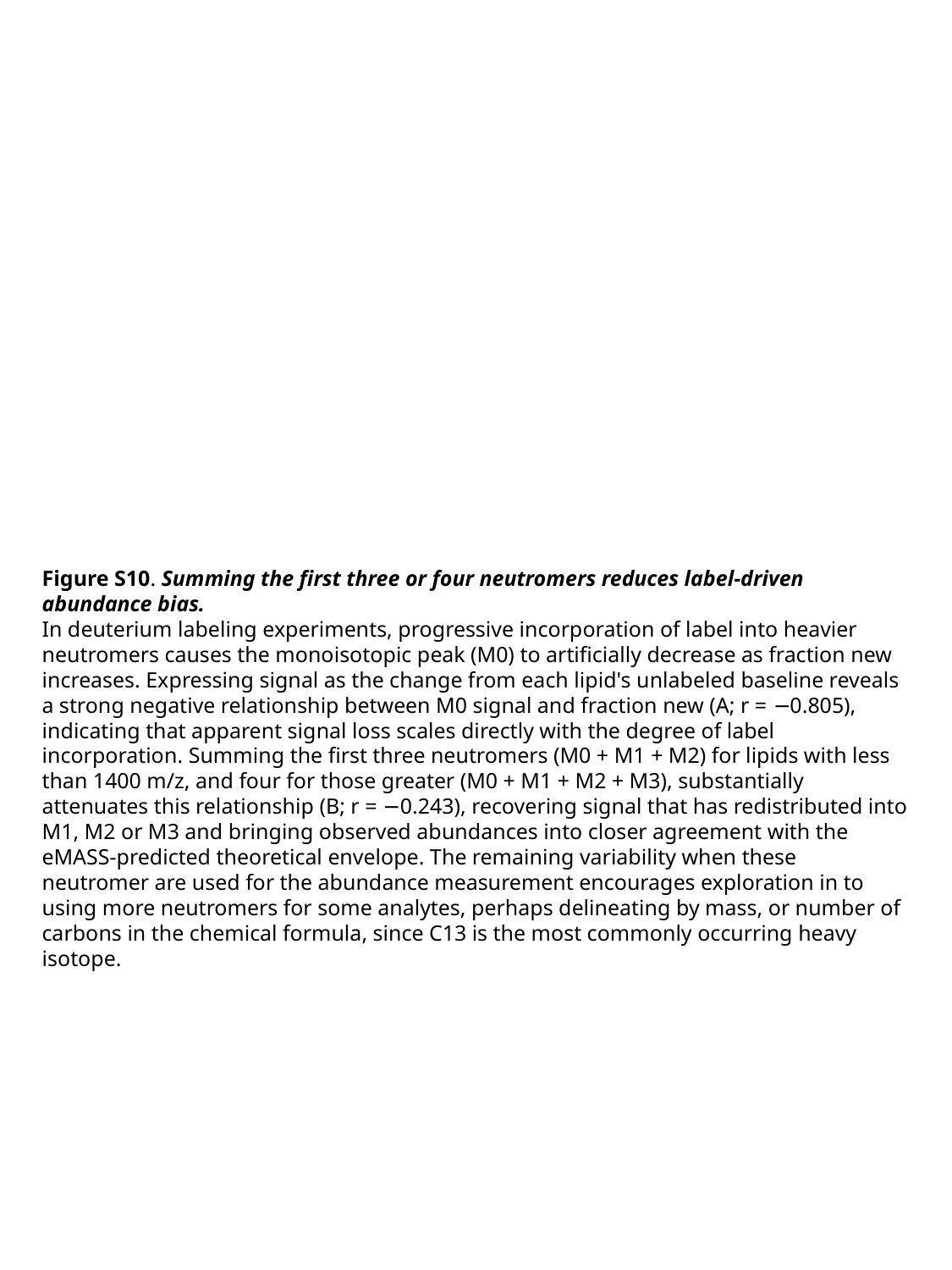

Figure S10. Summing the first three or four neutromers reduces label-driven abundance bias.
In deuterium labeling experiments, progressive incorporation of label into heavier neutromers causes the monoisotopic peak (M0) to artificially decrease as fraction new increases. Expressing signal as the change from each lipid's unlabeled baseline reveals a strong negative relationship between M0 signal and fraction new (A; r = −0.805), indicating that apparent signal loss scales directly with the degree of label incorporation. Summing the first three neutromers (M0 + M1 + M2) for lipids with less than 1400 m/z, and four for those greater (M0 + M1 + M2 + M3), substantially attenuates this relationship (B; r = −0.243), recovering signal that has redistributed into M1, M2 or M3 and bringing observed abundances into closer agreement with the eMASS-predicted theoretical envelope. The remaining variability when these neutromer are used for the abundance measurement encourages exploration in to using more neutromers for some analytes, perhaps delineating by mass, or number of carbons in the chemical formula, since C13 is the most commonly occurring heavy isotope.

### Slide 11
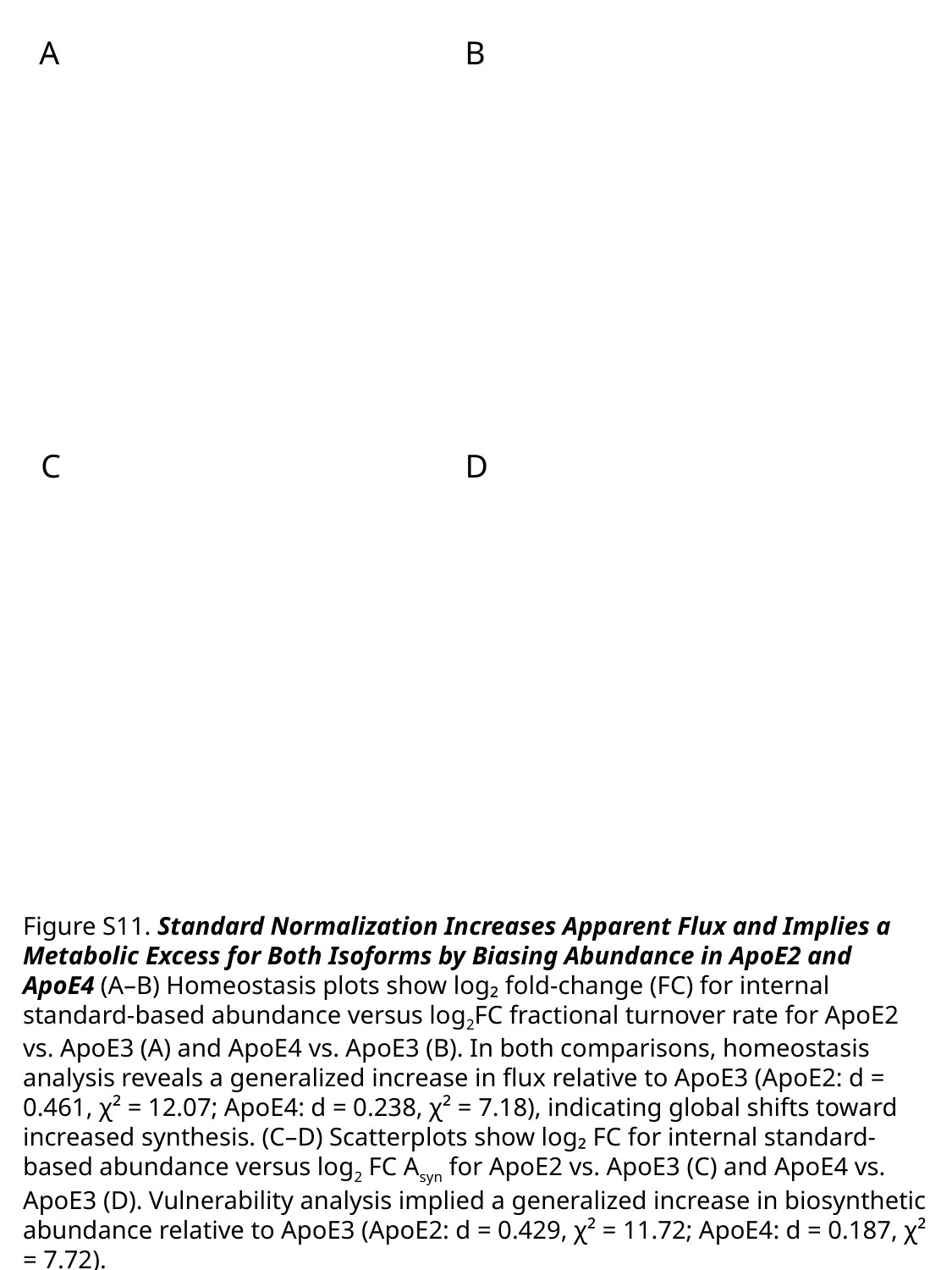

A
B
C
D
Figure S11. Standard Normalization Increases Apparent Flux and Implies a Metabolic Excess for Both Isoforms by Biasing Abundance in ApoE2 and ApoE4 (A–B) Homeostasis plots show log₂ fold‑change (FC) for internal standard-based abundance versus log_2_FC fractional turnover rate for ApoE2 vs. ApoE3 (A) and ApoE4 vs. ApoE3 (B). In both comparisons, homeostasis analysis reveals a generalized increase in flux relative to ApoE3 (ApoE2: d = 0.461, χ² = 12.07; ApoE4: d = 0.238, χ² = 7.18), indicating global shifts toward increased synthesis. (C–D) Scatterplots show log₂ FC for internal standard-based abundance versus log_2_ FC A_syn_ for ApoE2 vs. ApoE3 (C) and ApoE4 vs. ApoE3 (D). Vulnerability analysis implied a generalized increase in biosynthetic abundance relative to ApoE3 (ApoE2: d = 0.429, χ² = 11.72; ApoE4: d = 0.187, χ² = 7.72).

### Slide 12
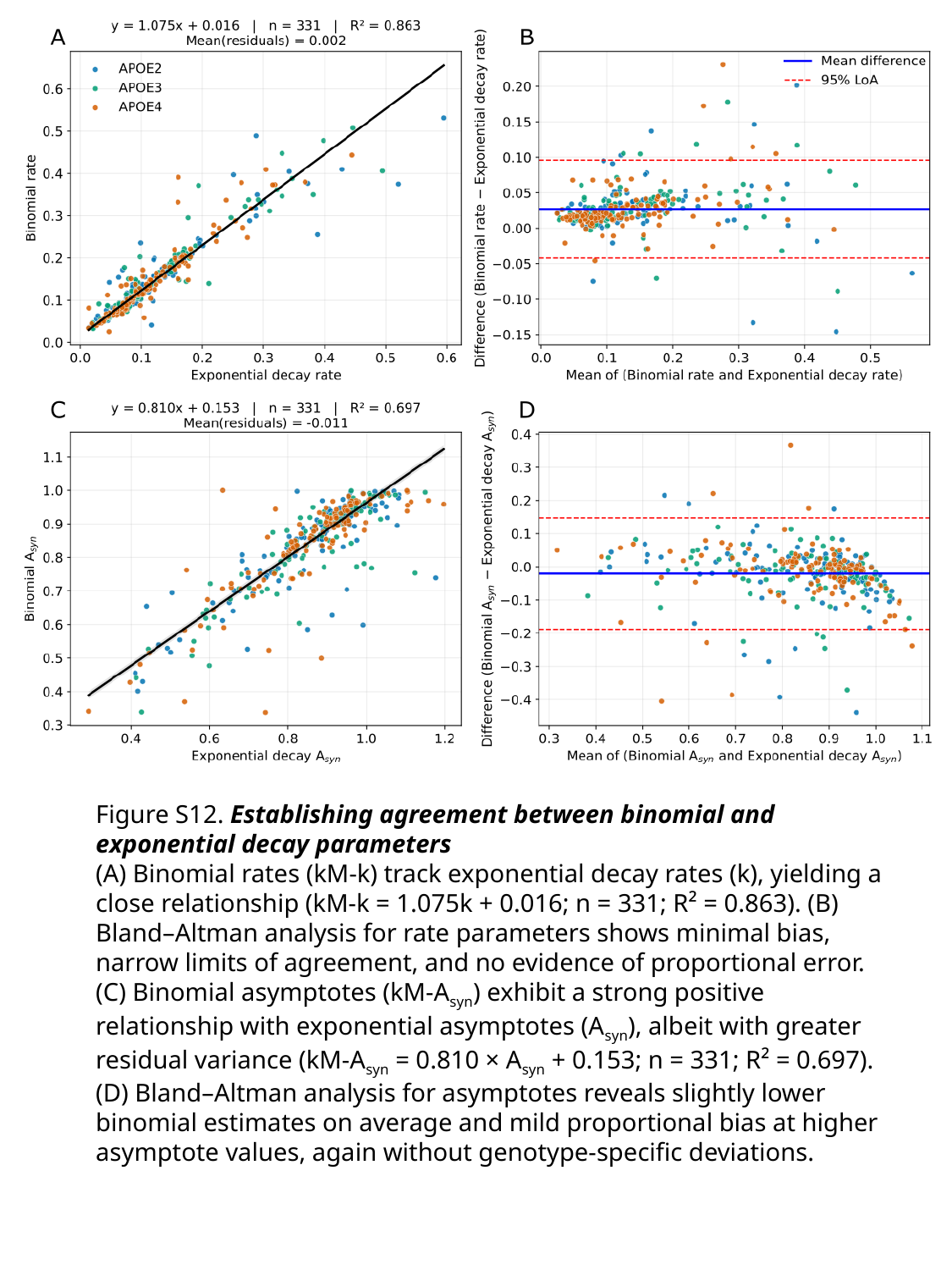

Figure S12. Establishing agreement between binomial and exponential decay parameters
(A) Binomial rates (kM‑k) track exponential decay rates (k), yielding a close relationship (kM‑k = 1.075k + 0.016; n = 331; R² = 0.863). (B) Bland–Altman analysis for rate parameters shows minimal bias, narrow limits of agreement, and no evidence of proportional error. (C) Binomial asymptotes (kM‑A_syn_) exhibit a strong positive relationship with exponential asymptotes (A_syn_), albeit with greater residual variance (kM‑A_syn_ = 0.810 × A_syn_ + 0.153; n = 331; R² = 0.697). (D) Bland–Altman analysis for asymptotes reveals slightly lower binomial estimates on average and mild proportional bias at higher asymptote values, again without genotype‑specific deviations.
